# DetectEV: a functional enzymatic assays to assess integrity and bioactivity of extracellular vesicles

**DOI:** 10.1101/2023.10.24.563745

**Authors:** Giorgia Adamo, Sabrina Picciotto, Paola Gargano, Angela Paterna, Samuele Raccosta, Estella Rao, Daniele Paolo Romancino, Giulio Ghersi, Mauro Manno, Monica Salamone, Antonella Bongiovanni

**Affiliations:** Cell-Tech HUB and Institute for Research and Biomedical Innovation (IRIB), National Research Council of Italy (CNR), Palermo 90146, Italy; Cell-Tech HUB and Institute of Biophysics (IBF) - CNR, Via Ugo La Malfa 153, 90146 Palermo, Italy; Department of Biological, Chemical and Pharmaceutical Sciences and Technologies (STEBICEF), University of Palermo, 90128 Palermo, Italy

**Keywords:** extracellular vesicles, integrity, bioactivity, functional enzymatic assay, quality check, standardization

## Abstract

The application of extracellular vesicles (EVs) as therapeutics or nanocarriers in cell-free therapies necessitates meticulous evaluations of different features, including their identity, bioactivity, batch-to-batch reproducibility, and stability. Given the inherent heterogeneity in EV preparations, this assessment demands sensitive functional assays to provide key quality control metrics, complementing established methods to ensure that EV preparations meet the required functionality and quality standards. Here, we introduce the detectEV assay, an enzymatic-based approach for assessing EV luminal cargo bioactivity and membrane integrity. This method is fast, cost-effective, and quantifiable through enzymatic units. Utilizing microalgae-derived EVs, known as nanoalgosomes, as model systems, we optimized the assay parameters and validated its sensitivity and specificity in quantifying the enzymatic activity of esterases within the EV lumen while also evaluating EV membrane integrity. Compared to conventional methods that assess physicochemical features of EVs, our single-step analysis efficiently detects batch-to-batch variations by evaluating changes in luminal cargo bioactivity and integrity across various EV samples, including differences under distinct storage conditions and following diverse isolation and exogenous loading methods, all using small sample sizes. The detectEV assay’s application to various human-derived EV types demonstrated its versatility and potential universality. Additionally, the assay effectively predicted EV functionality, such as the antioxidant activity of different nanoalgosome batches. Our findings underscore the detectEV assay’s utility in comprehensive characterization of EV functionality and integrity, enhancing batch-to-batch reproducibility and facilitating their therapeutic applications.

**Graphical Abstract:** 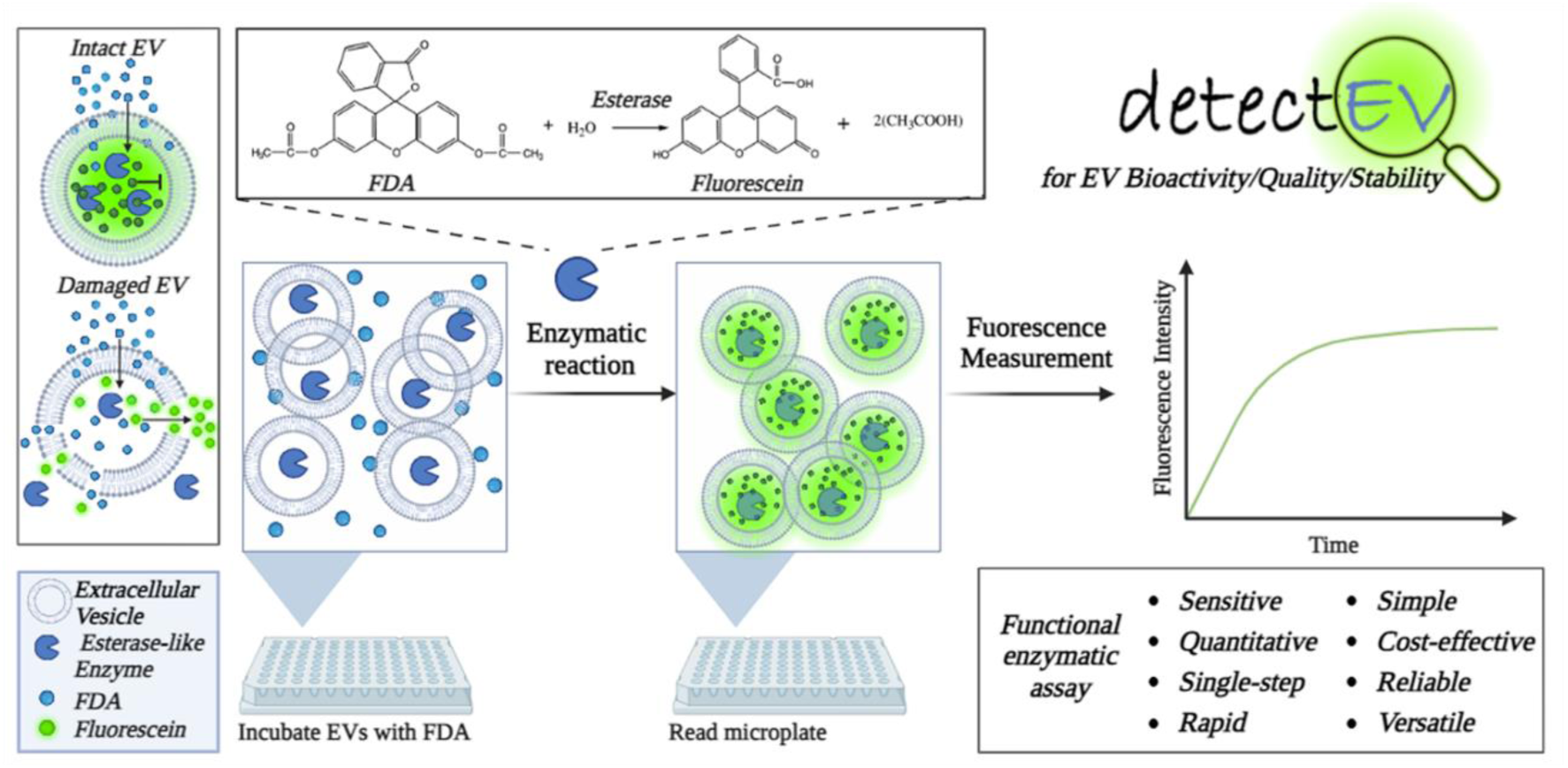

## Introduction

Extracellular vesicles (EVs) are essential signalling mediators involved in intercellular and inter-organism communications (Adamo et al., 2021; Raposo et al., 2013; Yáñez-Mó et al., 2015) . EV-based signaling depends on their lumen, membrane-integral or -associated cargo, which may be naturally-sourced (*e.g.*, bioactive molecules such as enzymes) or be exogenously loaded (*e.g.*, therapeutic molecules) (Chen Y, et al., 2021; O’Grady et al., 2022). In recent years, EVs have come to be considered one of the most promising innate effectors for cell-free therapy and bio-nanovehicles for drug delivery (Herrmann et al., 2021; Sun et al., 2021). However, despite the advances in the field, many challenges with EV-based therapies still stand (Herrmann et al., 2021). The therapeutic efficacy of EVs depends on their intrinsic properties, including the membrane stability (Russel et al., 2019). To validate the quality of EV preparations for therapeutic application and define the “Critical Quality Attributes” (CQA), EV-inherent features should be tested to qualify them as suitable for subsequent clinical functional testing applications (Yekula et al., 2020; Nguyen et al., 2020). In this view, to achieve an adequate assessment of EV bioactivity, appropriate functional assays need to be created to support common EV-related biophysical and biochemical quality controls (QCs).

Ideally, specific functional assays would predict whether a particular EV-preparation holds the potential to achieve its intended therapeutic effects or its “potency”, which could be further investigated in a “formal disease-specific” potency assays (Nguyen et al., 2020).

The MISEV-2023 guidelines and the EV-TRACK knowledgebase propose and support rigorous procedures required to document specific EV-associated functional activities to cope with the advance of EV research, assuring and improving the quality of these studies (Welsh et al., 2024; Van Deun et al., 2017). In particular, the MISEV-2023 recommendations for conducting functional studies encourage researchers to conduct dose-response and time-course studies, incorporate EV-negative controls, utilize EVs from different cell types for comparative analysis, and employ treated-EV samples to differentiate EV-specific effects from co-isolating materials. Moreover, it is suggested to explore the impact of EV separation or storage methods on EV activity to optimize experimental conditions (Welsh et al., 2024).

Furthermore, a functional assay should be simple and sensitive enough using a low amount of EV samples to facilitate the following studies (Pachler et al., 2017; Gimona et al., 2021). To date, only few *in vitro* and *in vivo* assays have been used to interrogate the potency of specific types of EVs; however, many of them failed for several challenges that make reliable assays difficult to set up (Nguyen et al., 2020; Ramirez et al., 2018). In the field of extracellular vesicles, it is expected that no single test can adequately measure all product attributes that can predict clinical efficacy. Further, the development of ubiquitous, standardisable and functional test for evaluating the bioactivity can be tricky for EVs because they are highly heterogeneous and show different bioactivity and biomolecular signatures (Ramirez et al., 2018). Despite this evidence, there is a need for a precise procedure to accurately qualify EV preparations, which can be applied to various types of EVs, regardless of their origin or preparation method (Gimona et al., 2021; LeClaire et.al 2021). It would be advisable to identify a common feature among vesicles that could predict EV functionality. An example could be evaluating the EV membrane integrity by measuring luminal enzymatic activity. In this context, a fascinating and unexplored aspect is the presence of a class of enzymes ubiquitously present in extracellular vesicles: esterases within EVs. Indeed, our search of the publicly available ExoCarta database revealed a noticeable and extremely robust association between esterase-like enzymes and vesicle-cargo proteins in EVs derived from many sources, including those of mammalian origin (mice, rats, and cows), several human body fluids (blood, saliva, and urine), and numerous human cancer and normal cell lines (more than 60 hits for “esterase” from the ExoCarta database, http://www.exocarta.org) (Gonzalez-Begne et al., 2009; Gonzales et al., 2009; Simpson et al., 2012). Furthermore, many recent proteomic analyses described the presence of esterase-like enzymes in EVs isolated from bacteria (like *Escherichia coli* and *Pseudomonas aeruginosa*), plants and fungi (*Candida albicans* and *Aspergillus fumigatus*) (Pocsfalvi et al., 2018; Rizzo et al., 2020; Choi et al., 2011; Garcia-Ceron et al., 2021; Bleackley et al., 2020; McMillan and Kuehn, 2022). A large number of enzymes, such as choline esterases, carboxylic ester hydrolases, lipases, and proteases, are member of the esterase’ superfamily, catalyzing the cleavage of ester bonds (Cygler et al., 1993). The measure of esterase activity can be performed by employing an enzymatic assay, using specific substrate, like the fluorescein diacetate. Fluorescein diacetate (*i.e.* 3’-6’-diacetyl-fluorescein, FDA) is a non-fluorescent fluorescein molecule conjugated to two acetate radicals (Adam and Duncan, 2001). This is a fluorogenic ester compound able to pass through phospholipid bilayers, like plasma and EV membranes. Indeed, when inside cells or EVs, fluorescein diacetate is hydrolyzed by non-specific esterases to produce a negatively charged membrane-impermeable green fluorescent molecule (*i.e.,* fluorescein) (Fontvieille et al., 1992). This membrane-permeable esterase substrate is commonly used as a probe to study microbial metabolic activity or to monitor the cell viability of fungi, plants, microalgae, and bacteria. It can be used both to measure enzymatic activity, essential for activating its fluorescence, and to evaluate membrane integrity, necessary for the intracellular or intra-vesicular retention of its fluorescent product (Ender et al., 2020; Gray et al., 2015).

In the EV-field, different approaches have described the use of membrane-permeant enzymatic substrates to label EVs, including carboxyfluorescein diacetate succinimidyl ester (CFSE), and calcein acetoxymethyl ester (calcein-AM) (Ender et al., 2020; Pospichalova et al., 2015; de Rond et al., 2018; Nikiforova et al., 2021; Tertel et al., 2022; Kormelink et al., 2016). Similarly to fluorescein diacetate, these molecules are cleaved by esterases, which are present inside cells as well as in secreted EVs (Gray et al., 2015). These experimental approaches have been developed to make EV fluorescent and detectable for flow cytometry applications or for fluorescent microscopy; however, these methods are not quantitative and do not directly provide information on EV-functional quality (Morales-Kastresana et al., 2017).

In this study, we introduced the detectEV assay as a new, simple, highly sensitive and precise functional test, able to define EV integrity and identity, by evaluating the EV-associated enzymatic activity, using a fluorescein diacetate-based enzymatic method (Sırt Çıplak and Akoğlu, 2020; Adam and Duncan, 2001). Throughout this assay, we considered various aspects crucial for working with EVs, as comparing the quality of different EV batches or EVs isolated using different methods (differential ultracentrifugation, dUC, versus tangential flow filtration, TFF). Additionally, we assessed the luminal enzymatic bioactivity and integrity of EVs stored under different conditions or manipulated using various physical loading techniques, using the proposed functional test and comparing these results with a standard method for EV evaluations, like nanoparticle tracking analysis (Welsh et al., 2024). Finally, we proved that the detectEV assay can predict the functionality of different preparations of EVs. The versatility of the detectEV assay for different type of EVs, including human ones, encouraged its application as functional enzymatic assay for rapid quality check (QC) of EVs, as in a single step analysis giving two important pieces of information: the bioactivity of a luminal molecules that is related to membrane integrity of EV preparations.

## Results

### Set up of the detectEV assay

To set up the detectEV assay, we used microalgae-derived extracellular vesicles (known as nanoalgosomes or algosomes) as small EV (sEV) models (Adamo et al., 2021; Picciotto et al., 2022). We isolated nanoalgosomes from the conditioned media of *Tetraselmis chuii* (*T. chuii*) microalgae, both in small and large scale using dUC and TFF methods, respectively. Next, we characterized them by evaluating selected MISEV-2023 quality parameters (Adamo et al., 2021; Welsh et al., 2024). These included biophysical characterization of size distribution and morphology by dynamic light scattering (DLS), nanoparticle tracking analysis (NTA) and atomic force microscopy (AFM), and a biochemical characterization of protein content using bicinchoninic acid assay (BCA assay) and immunoblot (IB) analysis for EV-biomarkers (Adamo et al., 2021; Paterna et al., 2022). Data relative to nanoalgosome characterization are reported in **Supporting Figure 1** (EV-TRACK ID: EV231004). In particular, nanoalgosomes exhibited a monodisperse size distribution of approximately 100 ± 10 nm, a rounded and homogeneous morphology, and the presence of enriched EV biomarkers such as Alix, H+-ATPase, and β-actin. Additionally, they are expected to demonstrate an appropriate EV particle number/protein ratio, in which 1 µg of total EV protein corresponding to a range of 5-10 x 10^9^ particles (Sverdlov, 2012).

**Figure 1.**
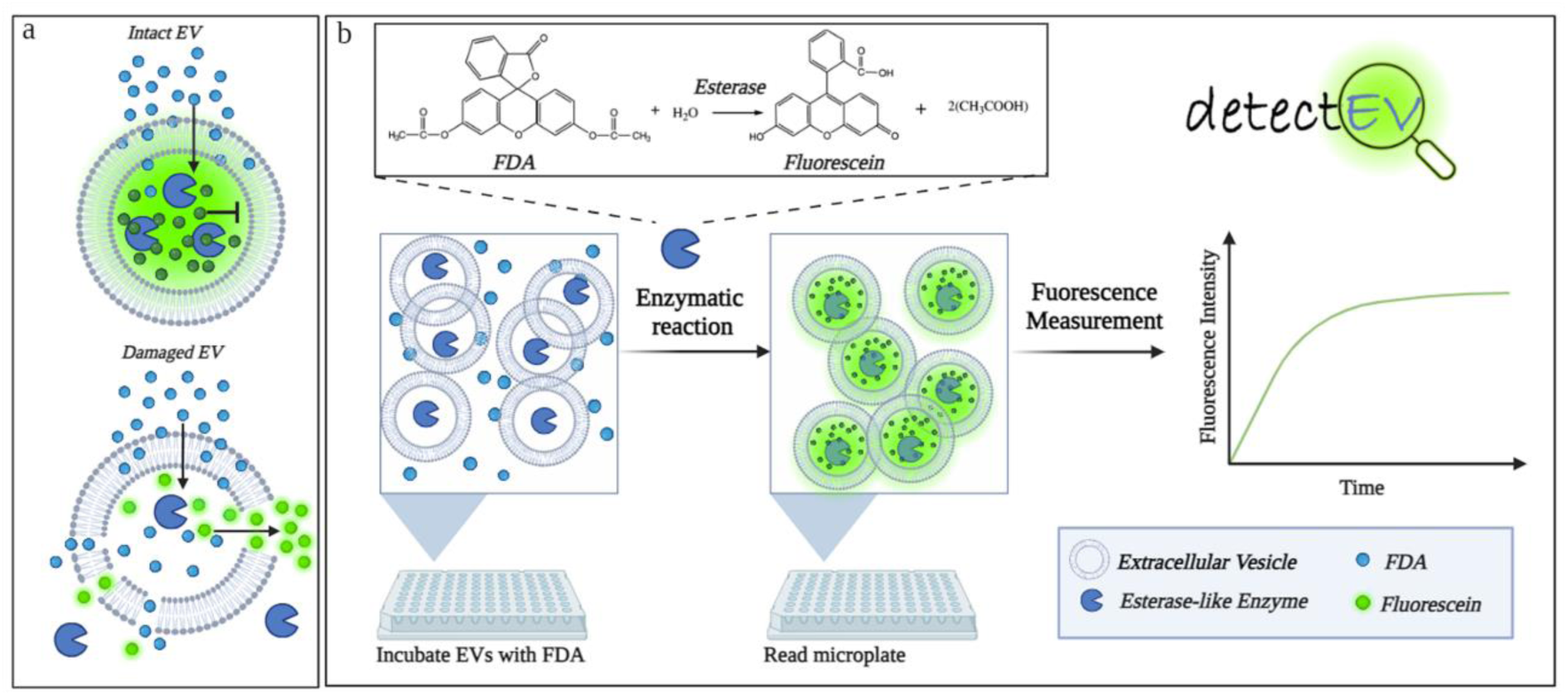
Schematic representation of the detectEV assay for evaluating the bioactivity and integrity of extracellular vesicles. a) Graphic representation of FDA feature in intact and damaged EVs. Intact EVs (top) retain their luminal contents, including esterase-like enzymes and the produced-fluorescein molecules, within the membrane, while damaged EVs (bottom) have compromised membranes leading to the leakage of their cargo contents. b) Overview of the detectEV assay procedure. EVs are incubated with fluorescein diacetate, which penetrates the vesicles. Inside intact EVs, esterase-like enzymes hydrolyze fluorescein diacetate to produce fluorescein, a fluorescent compound that is membrane impermeable. The enzymatic reaction’s mechanism is shown in the inset, indicating the hydrolysis of fluorescein diacetate to fluorescein and acetic acid. After the EV incubation with fluorescein diacetate substrate, the fluorescence intensity is measured using a microplate reader (excitation 488nm, emission 520nm). The fluorescence intensity over time is depicted in the graph on the right, showing the expected increase in fluorescence as the enzymatic reaction proceeds. Created with BioRender.com.

In the proof-of-concept of the proposed detectEV assay, the accumulation of fluorescein inside EVs is a measure of two independent parameters: enzymatic activity and membrane integrity (**Figure 1a)**. Therefore, in a first pilot test, we incubated a fixed amount of nanoalgosomes (2x10^10^) with fluorescein diacetate (FDA, 35 µmol) in 200µl of phosphate saline buffer solution (PBS) and checked for fluorescence emission with respect to EV-negative control (nanoalgosome vehicle solution, *i.e.,* PBS), using spectrofluorometer readings (**Figure 1b**). The fluorescence emission was detected over time, up to 16 hours (960 min). Interestingly, we got a rapid time-dependent increase of fluorescence intensity only in the nanoalgosome samples compared to the negative control. The trend of fluorescence increase over time exhibits a direct proportionality, *i.e.,* a linear relationship up to 8 hours, and subsequently reaches a plateau after approximately 15 hours, probably due to substrate depletion or enzyme inactivation (**Figure 2a**).

**Figure 2.**
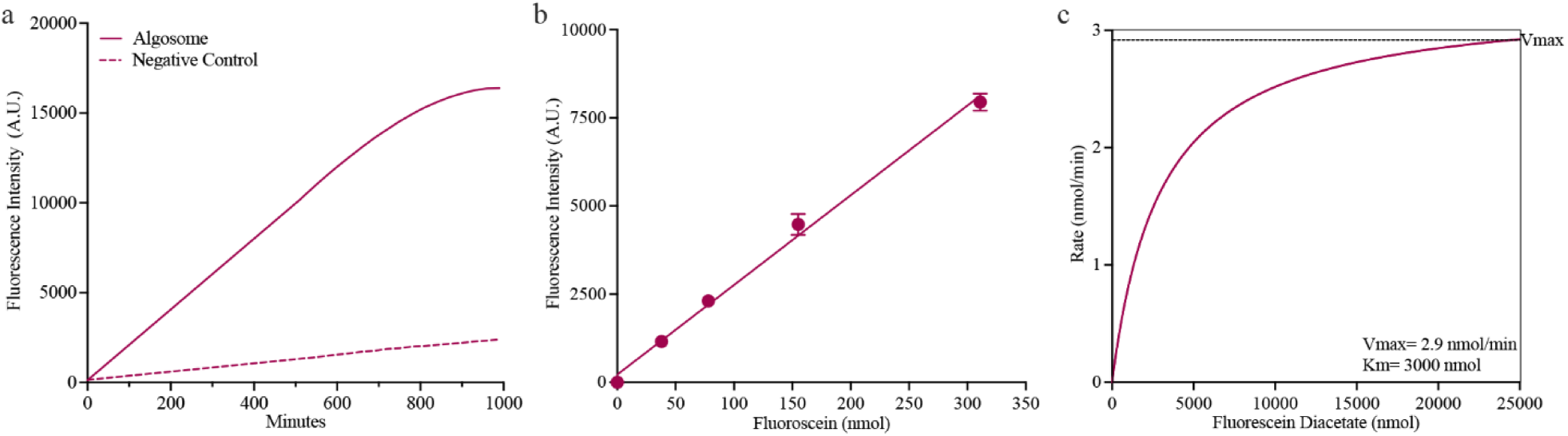
Set up and validation of the detectEV assay. a) Time-dependent increase in fluorescence intensity of microalgal-derived extracellular vesicles incubated with fluorescein diacetate. The fluorescence intensity expressed as arbitrary unit, A.U., (excitation 488nm, emission 520nm) of a representative microalgal-derived extracellular vesicles (*i.e.* nanoalgosomes or algosomes) sample incubated with fluorescein diacetate shows a significant increase over time, indicating the presence of esterase activity and intact vesicles. In contrast, the negative control (PBS, nanoalgosome vehicle control) shows minimal fluorescence intensity, confirming the specificity of the fluorescence signal to the enzymatic activity of the vesicles. b) Setting up of the detectEV assay. Calibration curve of fluorescein, depicting the linear relationship between fluorescence intensity and the concentration of fluorescein (nmol). Error bars represent the standard deviation of the mean fluorescence intensity of independent readings (n = 3). c) Michaelis-Menten graph illustrating the enzymatic cleavage rate of fluorescein diacetate (nmol/min) catalyzed by nanoalgosome enzymes. The curve demonstrates the relationship between the rate of reaction and the concentration of fluorescein diacetate (nmol). The maximum reaction rate (Vmax) is 2.9 nmol/min, and the Michaelis constant (Km) is 3000 nmol. These values could give an indication on the substrate concentration to be used to ensure that the enzymes are acting near Vmax (Bisswanger, 2014). Typically, a substrate concentration higher than the Km value is employed in enzymatic activity studies (Bisswanger, 2014). Based on this evidence, for establishing the right detectEV assay condition, we decided to fix the substrate concentration (*i.e.,*fluorescein diacetate) at 6 times the Km that is at 18 µmol (Michaelis et al., 2011; Duggleby, 1979).

Based on this result, fundamental parameters such as the temperature set at 22°C (*i.e.,* room temperature) and the duration of the assay limited to 3 hours (a time frame during which the reaction remains within the exponential phase and exhibits a linear trend) were defined. These parameters were established to ensure the test is rapid and sufficiently sensitive.

### Validation of the quantitative fluorescein diacetate-based functional enzymatic assay for EVs

For the validation of the analytical procedure, it is important to establish key parameters such as specificity, accuracy, and precision (Guidance for Industry Potency Tests for Cellular and Gene Therapy Products, 2011). Initially, we gated the appropriate approach to develop a functional assay for EVs that provides a universally quantifiable and standardizable read out. To accomplish that we had to determine the amount of fluorescein diacetate that is enzymatically cleaved during the assay, establishing a calibration curve to interpolate unknown quantities of cleaved fluorescein diacetate. Therefore, the fluorescein, corresponding to the reaction product of fluorescein diacetate hydrolysis, was deemed a reliable choice, and, in turn, it was generated a standard calibration curve using it (Dzionek et al., 2018). **Figure 2b** illustrates **the calibration curve** derived from serial dilutions of a fluorescein stock solution, where the fluorescence emission intensity exhibits a direct correlation with fluorescein concentration. The resulting calibration curve demonstrates a linear trend, facilitating the straightforward conversion of fluorescence intensity emitted following the enzymatic reactions (*i.e.,* fluorescein diacetate hydrolysis by active EV-related esterases) into nanomole (nmol) of produced fluorescein. Furthermore, in the development of the detectEV assay, we have chosen to adopt the enzymatic units (U) to quantitatively assess EV-related enzymatic activity. This because the specific enzymatic activity is universally expressed as U, representing the quantity of enzyme required to catalyse the conversion of one nanomole of substrate into product per minute (nmol/min), under the specified conditions of the assay method (Robinson, 2015).

Therefore, within the detectEV setup, enzymatic units (nmol/min) will be determined by utilizing a calibration curve of free fluorescein to calculate the total nmols generated over 180 minutes. Subsequently, total nmols will be divided by the overall time (180 minutes) to yield the final readout in nmol/min.

Additionally, in order to determine kinetic parameters of the enzymatic reaction, we performed an enzymatic kinetic study at increasing substrate concentrations, to measure the reaction rate (*i.e.,* the amount of substrate converted to product per unit of time) at constant nanoalgosome concentrations (2x10^10^ nanoalgosomes). This approach reflects the Michaelis-Menten model, a widely utilized method for studying enzyme kinetics (Bisswanger, 2014; Choi et al., 2017; Michaelis et al., 2011). As shown in **Figure 2c**, by plotting reaction rates against the respective substrate concentration, we obtained a canonical hyperbolic relationship, through which it was possible to derive kinetic parameters, including the maximum reaction velocity (Vmax, equal to 2.9 nmol/min) and the kinetic constant (as Michaelis constant, Km equal to 3 µmol) (Michaelis et al., 2011; Duggleby and Wood, 1989).

### Analytical sensitivity and specificity of detectEV assay

To establish the **analytical sensitivity** of the detectEV assay, we determined the limit of detection (LOD) by conducting a dose-response test to identify the lowest concentration of EVs that could be distinguished from the background noise. This background noise corresponds to the nanoalgosome buffer control (v/v of PBS solution) incubated with fluorescein diacetate.

Therefore, to determine the LOD, we incubated different nanoalgosome concentrations of the same preparation batch (ranging from 3x10^8^ to 2x10^10^ EVs) with 18 µmol of fluorescein diacetate. We found that the enzymatic assay is EV-dose responsive, with the minimum detection limit of 2x10^9^ nanoalgosomes (**Figure 3a**). Moreover, to assess potential influence of fluorescein diacetate auto-hydrolysis in different media, background fluorescence measurements were performed (**Supporting Figure 2a-b**). Fluorescein diacetate auto-hydrolysis was tested in microalgal F/2 medium and in human cell culture media (*i.e.* complete Dulbecco’s Modified Eagle Medium, DMEM with high and low glucose supplemented with EV-depleted serum), both in their original formulation and their post-purification form (using tangential flow filtration and differential ultracentrifugation). These controls ensured that any observed fluorescence was due to EV functionality rather than auto-hydrolysis of the fluorescein diacetate substrate. **Supporting Figure 2b** shows the background fluorescence measurements of the fluorescein diacetate up to 180 minutes: in F/2 medium, no significant auto-hydrolysis was observed, both in the original medium and the post-purification form. Conversely, both original formulation of DMEM media analysed (with high and low glucose) exhibited a high background signal, indicating significant fluorescein diacetate auto-hydrolysis, rendering the detectEV assay inapplicable for EVs in conditioned media like DMEM, as the high background could mask EV enzymatic activity-derived signals. While, fluorescein diacetate auto-hydrolysis of these post-isolation forms gave a signal comparable to the negative control (*i.e.*, PBS ranged from 400 to 800 arbitrary units of fluorescence), corroborating the robustness of the measurements. Consequently, all the detectEV-related experiments were conducted with EVs isolated and resuspended in PBS. Additionally, to verify whether the **analytical specificity** of the detectEV assay is associated with the bioactivity of active enzymes within intact EVs, we utilized nanoalgosomes treated with varying concentrations of a non-ionic detergent (Triton-X 100 at 0.1%, 0.25%, 0.5%, and 1%). This detergent permeabilizes lipid membrane bilayers and gently solubilizes proteins while maintaining their activity, including enzymatic activity (Perna et al., 2017; Jamur et al., 2010). As established in our prior study, complete nanoalgosome lysis was induced by treating with 1% Triton-X 100 (Adamo et al., 2021). Subsequently, we performed the detectEV assay using detergent-treated nanoalgosomes respect to not treated ones. The results showed in **Figure 3b** revealed higher enzymatic activity in the intact/no-treated nanoalgosomes, while this activity decreased with increasing doses of detergent, reaching no activity at the critical detergent concentration of 1%. This suggests that the measurable fluorescent signal relies on active esterases within the physiological environment of intact EVs. Identical results were observed when EVs were lysed, and proteins were denatured by boiling at 100°C for 10 minutes, thereby confirming the specificity of the proposed functional assay (**Figure 3b**).

**Figure 3.**
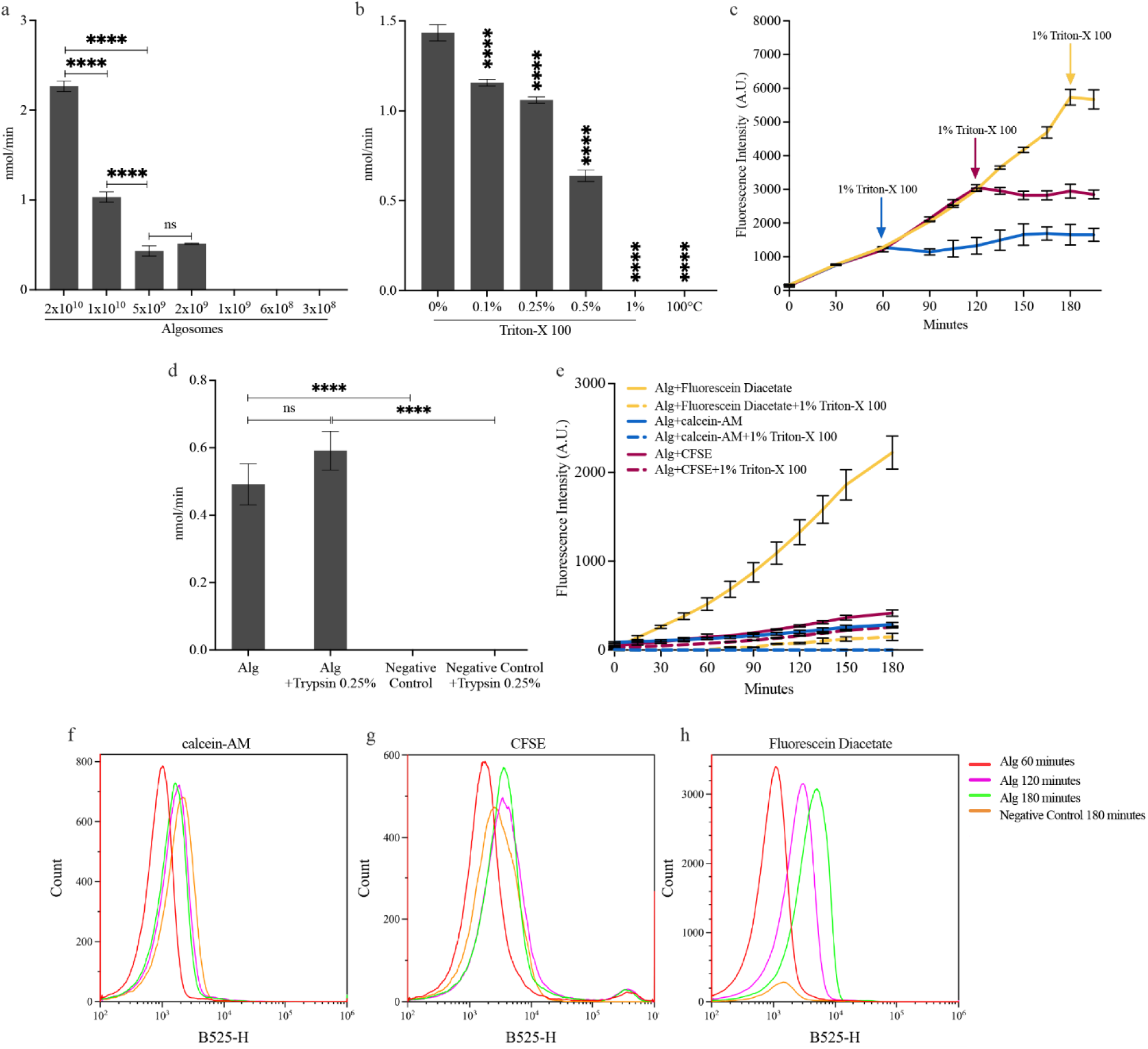
Evaluation of the detectEV enzymatic assay sensitivity and specificity, using nanoalgosome as EV-model. a) Detection limit determination using different concentrations of nanoalgosomes ranging from 2 x 10^10^ to 3 x 10^8^ sEVs. Error bars represent the standard deviation of the mean of independent experiments (n=3). One-way ANOVA was used to assess the statistical significance of the differences, showing ****p<0.0001 and non-significant differences (ns). b) Enzymatic activity comparison of untreated nanoalgosomes with those treated with various concentrations of Triton-X 100 detergent (0.1%, 0.25%, 0.5%, 1%) or boiled at 100°C. Error bars represent the standard deviation of the mean of independent experiments (n=3). One-way ANOVA was used to assess the statistical significance of differences between untreated nanoalgosomes and those treated with Triton-X 100 at 0.1%, 0.25%, 0.5%, 1%, as well as those boiled at 100°C, showing ****p<0.0001 for all conditions. c) Fluorescence intensity measurements of nanoalgosomes treated with 1% Triton-X 100 after 1 hour, 2 hours, and 3 hours following the addition of the fluorescein diacetate substrate. Error bars represent the standard deviation of the mean of the fluorescence intensity (subtracted from the respective PBS with or without 1% Triton-X 100 plus fluorescein diacetate background signal) of independent experiments (n=3). d) Comparison of the enzymatic activity of nanoalgosomes (Alg) and nanoalgosomes treated with 0.25% Trypsin, along with their respective negative controls (PBS without or with Trypsin 0.25%). Error bars represent the standard deviation (SD) of the mean from three independent experiments (n=3). One-way ANOVA was used to assess the statistical significance of differences between untreated nanoalgosomes and those treated with Trypsin 0.25%, showing non-significant differences (ns=not significant); the same test was applied to highlight the differences with the respective controls, showing ****p<0.0001. e) Comparative analysis of esterase-sensitive substrates using fluorescence-based plate reader. The graph show the fluorescence intensity expressed in arbitrary unit, A. U., (excitation 488nm, emission 520nm) up to 180 minutes for nanoalgosomes treated or not with 1% Triton-X 100 and incubated with calcein-AM, CFSE and fluorescein diacetate. The fluorescence intensity increases over time, indicating enzymatic activity only when the algosomes are incubated with fluorescein diacetate. Whereas for the calcein-AM and CFSE probes, only a slight increase in fluorescence was observed over time, reaching a very low A.U. even at 180 minutes. Error bars represent the standard deviation of the mean of the fluorescence intensity subtracted from the respective background signals measured in the corresponding negative controls, which are: PBS plus calcein-AM, CFSE, or fluorescein diacetate, with and without 1% Triton-X 100). Three independent experiments were considered (n=3). f-h) Comparative analysis of esterase-sensitive substrates using flow cytometry analyses. Histograms of counts versus fluorescence intensity (blue laser 488 nm) of nanoalgosome incubated 60, 120, and 180 minutes with (f) calcein-AM, (g) CFSE, (h) fluorescein diacetate. Negative control samples correspond to PBS incubated 180 minutes with calcein-AM, CFSE, or fluorescein diacetate.

Furthermore, to corroborate these results, we followed the enzymatic kinetics of nanoalgosomes treated with 1% Triton-X 100 after established time points (*i.e.,* after 1h, 2h, and 3h from the addition of the fluorescein diacetate substrate). As illustrated in **Figure 3c**, the fluorescence intensity was measured every 15 minutes during this time course test and, as expected, each no-treated sample exhibited a similar trend, namely a linear increase in fluorescence intensity over time. However, this increase ceased upon the addition of 1% Triton-X 100, remaining quite constant until the end of the assay. This trend demonstrates that the fluorescence intensity, corresponding to produced fluorescein, is correlated with membrane integrity of EVs. Indeed, when EVs are lysed the enzymatic reaction halts. Furthermore, the fluorescence values of fluorescein remain quite constant in intact vesicles compared to the respective lysed vesicles over time and for each condition, thereby validating the accurate application of the previously described calibration curve for free fluorescein, which is utilized in calculating nmols of fluorescein produced during the assay. Moreover, to corroborate the specificity of the detectEV assay, we performed a protease pre-treatment to verify that the quantified bioactivity is related to esterases located in the EV lumen. For this EV-protease treatment 0.25% trypsin was added to the EV samples and incubated at 37°C for 15 minutes (Chen C, et al., 2021),. By setting of the proper washing procedures, we successfully removed trypsin molecules from EV samples, checking also the proper nanoalgosome size distribution and concentration before and after these treatments (**Supporting Figure 2c-e**). As demonstrated in **Figure 3d**, the protease treatment of nanoalgosomes did not impact on esterase activity of protease-treated EV samples; thus, the quantified activity is attributed solely to the esterase within the lumen of the vesicles.

Based on the results from this section, the optimal conditions for the detectEV assay will involve incubating 2x10^10^ EVs with 18 µmol of fluorescein diacetate in a PBS solution, with a final reaction volume of 200 μL, in a 96-well plate. Enzymatic activity, measured through fluorescence emission (excitation at 488 nm, emission at 520 nm), will be followed using a spectrofluorometer for a duration of 3 hours at room temperature. The final readout corresponds to enzymatic units (nmol/min), which will be calculated using a calibration curve of free fluorescein to derive the total nmols produced during the assay. This value will then be divided by the total time (180 minutes) to obtain the final readout in nmol/min.

### Comparative analysis of esterase-sensitive substrates using flow cytometry and fluorescence-based microplate reader

We conducted a comprehensive comparison of the performance of fluorescein diacetate with other esterase-sensitive substrates, including CFSE and calcein-AM, and considered the lipophilic dye PKH67 as an esterase-sensitive negative control. This comparison was carried out using both flow cytometry and fluorescence-based microplate reader readings. The rationale behind this analysis was to explore the features of the detectEV assay by comparing it with similar methodologies and esterase-sensitive substrates. The experimental setup mirrored that of the detectEV assay, exploring the specificity using detergent treatment and recording the fluorescence background of each probe in PBS over time. As shown in **Figure 3e-h**, nanoalgosomes were incubated with these probes at concentrations reported for EV staining, and the samples were analysed both by fluorescence readings using microplate reader and flow cytometry and for 60, 120, and 180 minutes (Tertel et al., 2022; Ender et al., 2020). **Figure 3e** showed the results of the fluorescence intensity measured using a microplate reader over 180 minutes, maintaining conditions consistent with the detectEV assay. Calcein-AM and CFSE substrates incubated with nanoalgosomes showed a minor increase in fluorescence over time, and this increase was considerably smaller compared to EV-sample incubated with fluorescein diacetate, even using an equal number of 2x10^10^ EVs. As shown in **Supporting Figure 3a**, the results for PKH67-fluorescence intensity measurements in EV samples remained constant over time, as expected. Furthermore, calcein-AM and CFSE substrates exhibited a high fluorescent signal in the negative control (PBS) due to substrate auto-hydrolysis, resulting in elevated background values over time.

This phenomenon decreased their final fluorescent value, as the background had to be subtracted from those of the respective EV sample in the detectEV setting, thereby reducing the assay’s sensitivity when using these substrates compared to fluorescein diacetate (in which the background of the negative control showed a low fluorescence intensity after 180 minutes). Additionally, in detergent-treated EV samples, the CFSE substrate failed to accurately assess membrane integrity, as there was no difference in the fluorescence signal between samples treated with detergent and those not treated with detergent (**Figure 3e**). Accordingly, the flow cytometry approach showed that the fluorescence intensity increased over time only in EV samples incubated with the fluorescein diacetate substrate (**Figure 3h**). This increase was consistent with the esterase activity recorded using the detectEV approach, maintaining a low background fluorescence of the fluorescein diacetate in the negative control (PBS) up to 180 minutes. In contrast, the calcein-AM and CFSE substrates exhibited a slight increase in fluorescence intensity over time in EV samples (**Figure 3f-g**). However, the high background values of these substrates in the negative control hindered the evaluation of the EV specific enzymatic activity. For PKH67 staining, the fluorescence intensity remained constant over time in EV samples, reflecting its nature as a lipophilic probe that does not require enzymatic activation (**Supporting Figure 3b**).

These findings demonstrate that esterase-activated probes are ineffective in quantifying the esterase activity within EV lumen using flow cytometry. Although fluorescein diacetate produced the most significant increase in fluorescence, obtaining a quantitative measure of enzymatic activity through flow cytometry remains unfeasible when compared to the detectEV assay.

### detectEV assay to evaluate human EV integrity and functionality

To assess the detectEV assay’s applicability to other EVs, we examined its sensitivity, compatibility, and specificity using various EV types, including EVs isolated from human cell line conditioned media. Accordingly, we employed dUC to isolate sEVs from conditioned media of normal and tumoral mammary epithelial cells, and human embryonic kidney cells (1-7 HB2, MDA-MB 231 and HEK 293T cell lines, respectively). As for nanoalgosomes, we first characterized them by evaluating selected MISEV-2023 quality parameters, confirming the right size distribution, morphology and positivity to enriched EV biomarkers (data reported in **Supporting Figure 1**, EV-TRACK ID: EV231004) (Welsh et al., 2024). Following the biochemical and biophysical characterization of these three types of sEVs, we incubated 2 x 10¹⁰ HEK 293T derived sEVs with 18 µmol of fluorescein diacetate, using the same conditions as for the enzymatic activity evaluation of nanoalgosomes. As shown in **Figure 4a**, we observed a rapid, time-dependent increase in fluorescence intensity in the sEV samples compared to the negative control. The fluorescence increase follows a kinetic trend similar to that described for macroalgal-derived EVs.

**Figure 4.**
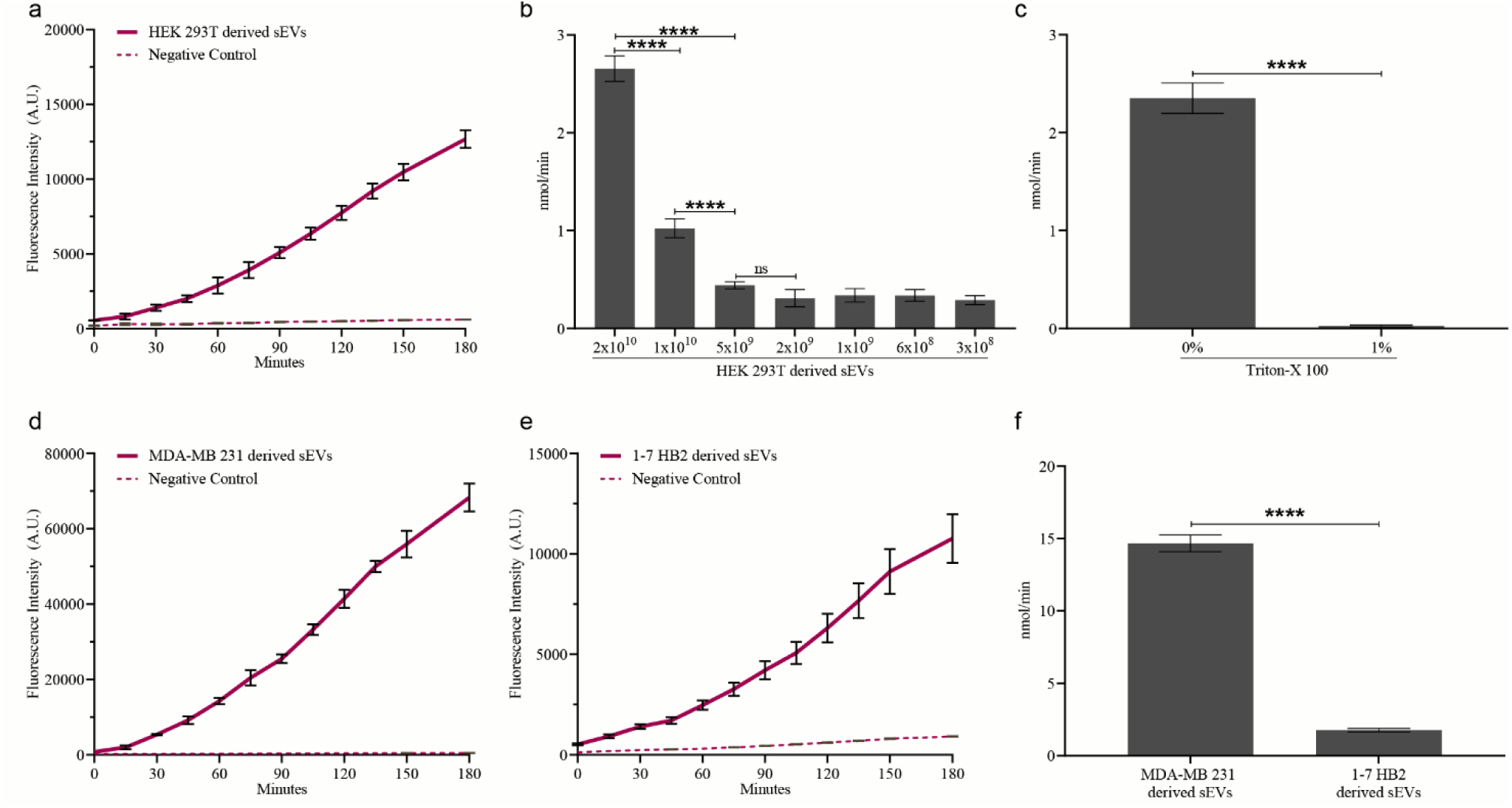
DetectEV assay on sEVs derived from different human cell lines. a) Time-dependent fluorescence increase of HEK 293T sEVs incubated with fluorescein diacetate. Fluorescence intensity (A.U., excitation 488nm, emission 520nm) of HEK 293T sEVs shows an increase over time, indicating esterase activity and intact vesicles. The PBS control shows minimal fluorescence, confirming signal specificity to vesicle enzymatic activity. b) Determination of enzymatic activity, expressed in nmol/min, using various concentrations of HEK 293T derived sEVs, ranging from 2 x 10¹⁰ to 3 x 10⁸ sEVs. Error bars represent the standard deviation of the mean of independent experiments (n=3). One-way ANOVA was used to assess the statistical significance of the differences, showing ****p<0.0001 and non-significant differences (ns=not significant). c) Enzymatic activity comparison of untreated HEK 293T derived sEVs with those treated with 1% Triton-X 100 detergent, expressed as nmol/min. Error bars represent the standard deviation of the mean of independent experiments (n=3). T-test was used to assess the statistical significance of differences between untreated nanoalgosomes and those treated with Triton-X 100 at 1%, showing ****p<0.0001. d-e) Time-dependent fluorescence increase of sEVs isolated from MDA-MB 231 (breast cancer cells), 1-7 HB2 (normal mammary epithelial cells), respectively. sEVs were incubated with fluorescein diacetate, and their fluorescence intensity (A.U., excitation at 488 nm, emission at 520 nm) increased over time, indicating esterase activity and the presence of intact vesicles. In contrast, the PBS control exhibited minimal fluorescence, confirming that the signal is specific to the vesicles’ enzymatic activity. f) Enzymatic activity (nmol/min) of sEVs isolated from MDA-MB 231 and 1-7 HB2 cells. Error bars represent the standard deviation of the mean of independent experiments (n=3). T-test was used to assess the statistical significance of differences between MDA-MB 231 vs 1-7 HB2 derived sEVs, showing ****p<0.0001.

Next, we used different concentrations of HEK 293T derived sEVs to determine the LOD by conducting a dose-response test, as previously described for nanoalgosomes. We incubated different concentrations of HEK 293T derived sEVs from the same preparation batch (ranging from 3 x 10⁸ to 2 x 10¹⁰ EVs) with fluorescein diacetate. Similar to nanoalgosomes, we found that the enzymatic assay is dose-responsive, with a specific signal up to 1 x10^10^ HEK 293T derived sEVs (**Figure 4b**). We then evaluated the specificity of the detectEV assay using detergent-treated and untreated HEK 293T derived sEVs. The results, shown in the **Figure 4c**, revealed enzymatic activity of approximately 2.2 nmol/min in 2 x 10¹⁰ intact/untreated vesicles, while no activity was detected following 1% Triton-X 100 treatment.

Subsequently, we incubated 2 x 10¹⁰ of MDA-MB 231 and 1-7 HB2 derived sEVs with 18 µmol of fluorescein diacetate, following the detectEV assay setup described earlier. The results in **Figure 4d-f** confirmed the presence of reliable enzymatic-dependent activity for both of EV types analyzed, demonstrating detectEV assay’s compatibility with human EVs. Interestingly, breast tumor-derived sEVs showed significantly higher enzymatic activity (five-fold increase) compared to sEVs isolated from normal breast cells, opening new perspectives for detectEV assay’s application in tumor diagnosis (**Figure 4f**).

### Application of detectEV assay

#### Comparison between EV-isolation methods

Depending on experimental settings and/or the production scale, establishing the most efficient method for EV separation while ensuring sample quality is essential; we previously used various procedures (*e.g.*, dUC and TFF) to efficiently isolate nanoalgosomes from microalgal conditioned media (Adamo et al., 2021; Picciotto et al., 2021). Here, we compared the quality of nanoalgosomes from the same *T. chuii* culture batch, isolated in parallel by dUC and TFF, applying the detectEV assay. The results indicated that nanoalgosomes isolated by either method showed the same bioactivity/stability, confirming the good quality of both nanoalgosome preparations (**Figure 5a**).

**Figure 5.**
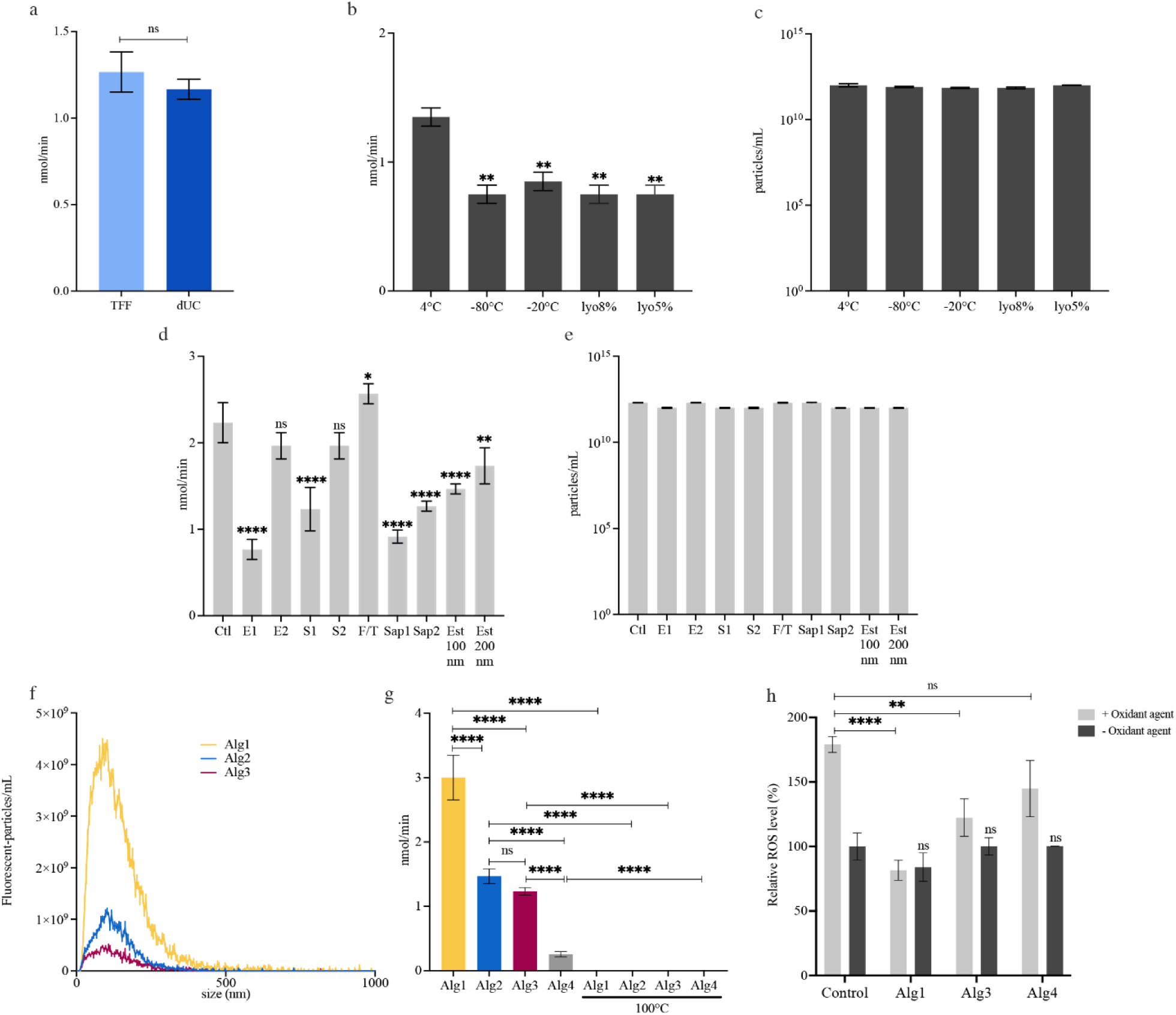
DetectEV assay applications. a) Comparison between nanoalgosomes isolated with different methods. Enzymatic activity (nmol/min) of nanoalgosomes isolated by tangential flow filtration (TFF) and differential ultracentrifugation (dUC). The error bars represent the standard deviation of the mean of independent experiments (n=3). Statistical significance was determined using one-way ANOVA: TFF vs dUC, ns=not significant. b-c) Comparison between nanoalgosomes stored under different conditions. Effect of different storage conditions on nanoalgosomes (*i.e.,* storage for 10 days at 4°C, -20°C, -80°C, and lyophilized with 5% and 8% sucrose) analyzed by (b) the detectEV assay (nmol/min) and by (c) NTA (particles/mL). In (b) error bars represent the standard deviation of the mean of independent experiments (n=3); in (c) distributions errors are calculated on 5 replica for each conditions of independent experiments (n= 3). One-way ANOVA was used to assess the statistical significance of the differences: 4°C vs -80°C, -20°C, lyo8%, lyo5%; **p<0.01; no significant differences were observed in the NTA data. d-e) Comparison between nanoalgosomes subjected to various loading methods. Effect of different loading methods on nanoalgosomes analyzed by (d) detectEV assay (nmol/min) and (e) NTA (particles/mL). Electroporation settings included E1 (125 μF, 400 V, 2 pulses of 20 ms) and E2 (125 μF, 250 V, 2 pulses of 30 ms). Sonication settings included S1 (ultrasonic probe, 20% amplitude, six cycles of 30 s on/off, total 3 min, 2 min cooling, 60 min incubation at 37°C) and S2 (ultrasonic bath, 40 KHz, 40% amplitude, two cycles of 30 s on/off, 60 min incubation at 37°C). F/T consist in three times freeze-thaw cycles at -80°C for 30 min and at RT for 30 min. Saponin treatment included two conditions: Sap1 (0.1 mg/mL) and Sap2 (0.002 mg/mL). Extrusion used polycarbonate membrane filters of 100 nm (Est100nm) and 200 nm (Est200nm) pore size, with each sample extruded 31 times. In (d) error bars represent the standard deviation of the mean of independent experiments (n=3). In (e) distributions errors are calculated on 5 replica of independent experiments (n=3). One-way ANOVA was used to assess statistical significance: Ctl vs (E1, S1, Sap1, Sap2, Est100nm), ****p<0.0001; Ctl vs (E2, S2), ns=not significant; Ctl vs F/T, *p<0.05; Ctl vs Est200nm, **p<0.01; no significant differences were observed in the NTA data. f-g) Comparison between different nanoalgosome batches. f) Size distribution of different nanoalgosome batches (Alg1, Alg2, Alg3) stained with a green lypophilic dye (Di-8-ANEPPS), measured by fluorescent-nanoparticle tracking analisys (F-NTA, using NanoSight NS300, with a 500LP filter and a laser wavelength of 488 nm); distributions errors are calculated on 5 replica of the same batch; F-NTA measurement for Alg4 was unreliable because below the detection limit. g) detectEV assay on different nanoalgosome batches (Alg1, Alg2, Alg3, Alg4). Error bars represent the standard deviation of the mean of 3 replica of the same batch. One-way ANOVA was used to assess the statistical significance of the differences where indicated, showing non-significant differences (ns=not significant); ****p <0.0001. h) Antioxidant activity of different nanoalgosome batches in 1-7 HB2 cells. ROS production in 1-7 HB2 cells treated with 0.5 µg/mL nanoalgosome for 24h, with/without oxidant agent (250 µM TBH), normalized to negative control (untreated cells). Values are presented as means ± standard deviation from three independent experiments (n=3). Statistical significance of the differences was assessed using one-way ANOVA: 1-7 HB2 (-Oxidant agent) vs 1-7 HB2 treated with Alg1, Alg3, Alg4 (-Oxidant agent), ns=not significant; 1-7 HB2 (+ Oxidant agent) vs 1-7 HB2 treated with Alg1 (+ Oxidant agent), ****p<0.0001; 1-7 HB2 (+ Oxidant agent) vs 1-7 HB2 treated with Alg3 (+ Oxidant agent); **p<0.01; 1-7 HB2 (+ Oxidant agent) vs 1-7 HB2 treated with Alg4 (+ Oxidant agent), ns=not significant.

#### Storage conditions

We applied the detectEV assay to monitor the effects of one-week storage under different conditions (including 4°C, -20°C, -80°C, lyophilization in 5% and 8% sucrose) on nanoalgosome bioactivity/stability (**Figure 5b-c)**. In parallel, we performed NTA readings to monitor the size distribution and concentration of nanoalgosomes stored under these conditions. For all the samples analysed, we did not observe changes in size distribution with an average size of 100±20 nm and a quite similar nanoparticle concentration between the samples (**Figure 5c, Supporting Figure 4**). Conversely, as reported in **Figure 5b**, the detectEV assay highlighted a significant and quantifiable reduction of enzymatic activity in EV, less than 1 nmol/min in frozen (-20°C or -80°C) or lyophilized EV-samples, compared to those stored at 4°C for a week. These results demonstrate that the assay can efficiently identify the most appropriate storage method for EV.

#### Loading methods

EVs are considered a promising novel drug delivery system. To this end, we explored the most reported loading methods for encapsulating biotherapeutics into EVs to determine how external stimuli, such as electrical (i.e., electroporation) or ultrasonic (i.e., sonication) interventions, affect vesicle membrane integrity. This evaluation was conducted without introducing an exogenous cargo, focusing instead on assessing the effectiveness of the proposed assay in detecting changes related to EV integrity. For each method, we used the most commonly reported EV loading settings before applying the detectEV assay (Chen C, et al., 2021; Herrmann et al., 2021) (**Figure 5d)**. Contextually, we monitored changes in nanoalgosomes size distribution and concentration after the application of each loading method, using NTA (**Figure 5e**). In **Figure 5d**, we reported detectEV results, which highlight that freeze and thaw approach appear to be the “gentlest” loading method, with less impact on nanoalgosome features. In contrast, for some conditions used such as electroporation (E1), sonication (S1), saponification (Sap1-2), or extrusion, we observed a significant decrease in EV-enzymatic activity below 1 nmol/min, probably due to harsh membrane perturbation or loss of vesicle integrity. Interestingly, this difference could not be detected by NTA analysis, as the size (inside the range of 100±20 nm for all samples) and concentration of these samples were similar with the untreated-control (**Figure 5e, Supporting Figure 5**). The successful application of the assay for evaluating EVs during loading could be the starting point for developing effective EV-based therapeutics.

#### Batch-to-batch reproducibility

To assess the detectEV assay’s capability in evaluating batch-to-batch reproducibility, we selected four nanoalgosome batches (named Alg1, Alg2, Alg3, Alg4) that exhibited variations in quality during the QC analyses. Following the separation of nanoalgosomes through tangential flow filtration, the application of EV-based QC methods for each production allowed us to identify differences in terms of quality, primarily attributed to the performance or half-life of the TFF cartridges (Adamo et al., 2021; Paolini et al., 2022; Paterna et al., 2022). These differences was revealed by different EV analyses, including fluorescent-NTA readings, after staining 10^10^ nanoalgosomes with di-8-butyl-amino-naphthyl-ethylene-pyridinium-propyl-sulfonate (Di-8-ANEPPS, excitation 488 nm, emission 630 nm), a specific lipophilic fluorescent dye that emit a green fluorescence when it is bound to lipid bilayer. For this reason this dye could help to discriminate between EVs and non-vesicle co-isolates (Adamo et al., 2021). As reported in **Figure 5f**, the F-NTA results showed that the selected four batches had a quite similar size distribution, but differences in terms of percentage of Di-8-ANEPPS-positive EVs with respect to the total EVs tracked with NTA in scattering. More specifically, Alg1=10%; Alg2=3%; Alg3=2%, Alg4= unreliable measurement.

Subsequently, we applied the detectEV assay for these EV-preparations, using an equal amount of vesicles for all batches (2x10^10^ nanoalgosomes). The results shown in **Figure 5g** identified Alg1 as the one with high enzymatic activity compared to the other three batches analysed, and are consistent with the F-NTA results (**Figure 5f**). All these results suggest that the proposed functional enzymatic assay allows for direct and quantitative comparison of EV quality across different isolation methods, storage conditions, and loading conditions. Additionally, it enables the validation of EV preparations and monitoring of batch-to-batch variability during EV production.

### detectEV prediction of EV functionality

To showcase the predictive capability of the detectEV assay in evaluating the functionality of EVs, we performed an antioxidant activity test as a bioactivity assessment, given the robust antioxidant properties of nanoalgosomes (Adamo et al., 2024). We selected nanoalgosome batches with significant differences in quality, as indicated by the detectEV results showed in **Figure 5g**. Specifically, we chose Alg1 and Alg3 batches that exhibited significantly higher esterase activity compared to the Alg4 batch.

Subsequently, 1-7 HB2 cells were exposed to 0.5 μg/mL of Alg1, Alg3, and Alg4 (approximately 10^10^ EVs) to assess their antioxidant bioactivity, particularly in terms of countering reactive oxygen species (ROS) production in cells stressed with the oxidant agent. The results demonstrated that Alg1 and Alg3 effectively mitigated the oxidative stress induced by the oxidant agent, restoring ROS levels near to physiological values, with Alg1 exhibiting strong antioxidant activity (**Figure 5h**). In contrast, Alg4 failed to alleviate the stress, indicating no significant antioxidant activity. These outcomes align seamlessly with the detectEV results, highlighting its ability to predict the functionality of various EV preparations.

## Discussion

The field of extracellular vesicles has made significant advancements; yet determined challenges persist, primarily centred around the lack of reliable functional assays for assessing EV bioactivity (Nguyen et al., 2020). The intricate nature of EV heterogeneity and their regulated mechanisms across different diseases adds complexity to developing robust and universal functional assays. Despite the diversity of available potency assays, challenges in quantitativeness, sensitivity, accuracy, precision, and robustness persist, emphasizing the urgent need for refining and establishing specific test capable of predicting EV functionality effectively and for different applications (Pachler et al., 2017; Gimona et al., 2021; Nguyen et al., 2020; Ramirez et al., 2018). Also, given the complex nature of EV-based therapeutics, multiple assays are often necessary to capture the various mechanisms of action (MoAs) EV may exhibit, ensuring compliance with regulatory requirements for efficacy and safety. These comprehensive functional evaluations are vital for the successful and safe development of EV-based therapeutic solutions. The bioactivity of EVs can be driven by surface signaling mechanisms, through the transfer of their internal cargo to recipient cells, or a combination of both. In surface signaling, EVs may interact with target cells via receptor-ligand binding or other membrane interactions. On the other hand, cargo transfer refers to the process in which EVs deliver their internal content directly into recipient cells, influencing cellular functions from within.

In this context, this study explores the potential to leverage the presence of active cargo enzymes within EVs for the development of an enzymatic assay to assess the EV integrity. Specifically, our focus has been on esterase-like enzymes in EVs, as their presence is reported in different proteomic datasets, showing associations with EVs from various sources (Gonzales et al., 2009; Gonzalez-Begne et al., 2009; Pocsfalvi et al., 2018; Rizzo et al., 2020; Choi et al., 2011; McMillan and Kuehn, 2022). The proposed detectEV assay specifically aims to quantitatively measure esterase activity within EVs, offering a single-step analysis for assessing the quality of EV preparations, using small-sized samples. Notably, this assay does not require washing steps, making it a faster and cost-effective method that offers several advantages, including simplicity. The establishment of the detectEV assay involved a meticulous validation of specific parameters (*e.g.*, EV and fluorescein diacetate concentrations, reaction volume, reaction buffer, time, and temperature), ensuring analytical sensitivity and specificity, in line with MISEV-2023 guidelines (Welsh et al., 2024). Nanoalgosomes, which are microalgal derived-EVs, were employed as EV models in this validation process. While our results demonstrate that the established experimental settings efficiently measure the enzymatic activity of nanoalgosomes, as well as of EVs derived from different mammalian cell lines, these parameters could be further adapted ad hoc for EVs from various origins or types, like large EVs or oncosomes. Indeed, different types of EVs could exhibit distinct stability or carry different combinations of esterase-like enzymes as cargo, possibly with high enzymatic activity. Consequently, it might be possible to slightly adapt the assay condition, such as by decreasing the amount of fluorescein diacetate, adjusting the reaction time, or vesicle concentration, to establish minimum and maximum ranges of enzymatic activity that qualify vesicle preparations for subsequent analyses. Therefore, the versatile applicability of the detectEV assay across diverse cellular sources allow for its extension to every EV types with esterase-like activity. This positioning makes it a potentially universal measure of EV functionality. The innovation introduced by the detectEV assay is its ability to simultaneously measure both the esterase’s enzymatic activity and membrane integrity of vesicles, which is crucial for EV functionality. Specifically, our results illustrated that when the vesicle integrity is perturbed and/or completely lost, the enzymatic activity decrease or becomes undetectable by the detectEV assay. This observation could be attributed to several reasons. First of all, there could be a potential reduction in the enzymatic activity of esterases when released into the extra-vesicular environment, as compared to the optimal condition of the vesicular lumen. The protease treatment of EVs does not affect esterase activities as they are indeed located in the lumen of EVs, demonstrating that detectEV give a measure of bioactive enzymes inside intact vesicles. Moreover, we demonstrated that the detectEV assay extends similar approaches using FDA-like molecules (*e.g.*, calcein-AM or CFSE) already employed to label EVs for flow cytometry analysis (Ender et al., 2020; Nikiforova et al., 2021; Tertel et al., 2022; Kormelink et al., 2016). Our comparative studies showed that using fluorescein diacetate for functional evaluations of EVs outperformed other esterase-sensitive substrates in both flow cytometry and microplate reader analyses, confirming its superiority for assessing the bioactivity of EV luminal cargo and membrane integrity. To note, a precise readout value of EV bioactivity expressed as enzymatic units (nmol/min) is a unique feature that enhances the novelty of the detectEV method, as compared to the fluorescence intensity values (expressed in arbitrary units) provided by calcein-AM or CFSE-based approaches, which allow for a semi-quantitative final value.

Another relevant aspect to consider is the detectEV assay’s versatility for different applications. One of these includes monitoring storage conditions of EV preparations. Various studies have demonstrated how different storage conditions can affect EV characteristics, including membrane stability and potency (Sivanantham and Jin, 2022; Jeyaram and Jay, 2017; van de Wakker et al., 2022). Recent comprehensive studies have compared different storage strategies to identify appropriate conditions for stabilizing EV preparations, especially for therapeutic applications (Görgens et al., 2022, Kusuma et al., 2018; Lener et al., 2015; Lőrincz et al., 2014;). Following EV sample storage, they are typically analyzed using methods like NTA, or flow cytometry (Sokolova et al., 2011; Lőrincz et al., 2014). Our results show that the detectEV assay is more effective than NTA approaches in highlighting slight differences in EV functionality after different storage methods. Numerous studies have highlighted the advantages of using EVs as drug delivery systems in preclinical models due to their low toxicity, high targeting capacity, and slow clearance (Gangadaran and Ahn, 2020). Exogenous cargo can be loaded into EVs using various physical methods, including electroporation, sonication, saponin-assisted loading, freeze-thaw cycles, and extrusion (Fu et al., 2020; Chen C, et al., 2021; Van Deun et al., 2020) . However, these methods may compromise EV functionality or damage the integrity of their membranes, although the phospholipid bilayer typically restores its integrity quickly after membrane perturbations (Han et al., 2021; Rankin-Turner et al., 2021). In this context, we have demonstrated that the detectEV assay can assess the effects of different loading methods on EVs, highlighting changes or alterations in their luminal bioactivity post-loading; this is because of the assay closely correlates with the bioactivity and quality of the vesicles, as well as the integrity of their membranes. Such capability is crucial for establishing and validating optimal exogenous loading strategies for specific types of EVs, particularly those carrying cargo lacking active components that can react with fluorescein diacetate, thus facilitating their application as nanocarriers for drug delivery in therapeutic contexts (Fu et al., 2020; Lener et al., 2015). Additionally, our findings suggest that the detectEV assay can determine the most effective methods for monitoring specific experimental settings. According to MISEV-2023 guidelines, the quality of EVs can vary significantly based on their source and the scale of production, which necessitates different isolation procedures (Welsh et al., 2024). Our results demonstrate that another valuable application of the detectEV assay is in determining the most suitable isolation methods to monitor specific experimental settings, ensuring the preparation of high-quality bioactive vesicles. Another application described here is its ability to highlight batch-to-batch variation between EV preparations. Indeed, the detectEV results were in complete agreement with the F-NTA data obtained from diverse nanoalgosome batches, showing that higher quality vesicles (Alg1) had also higher enzymatic activity. Additionally, the nanoalgosome batch Alg1 has been shown to be the nanoalgosome preparation that most efficiently counterbalances cell-oxidative stress *in vitro*, as observed in a specific functional test for nanoalgosomes (antioxidant activity assay). This suggests that the detectEV result could predict which vesicle preparation possesses greater specific functionality for subsequent potency test. The assay’s future potential to be explored involve developing it as a diagnostic tool to identify pathological signs, such as tumor-derived EVs, based on the evaluation of EV enzymatic activities in liquid biopsy samples (*e.g.*, plasma and urine), aligning with the growing importance of EVs in theranostics (Liang et al., 2021). To conclude, we propose that alongside efforts to harmonize EV nomenclature and characterization, the detectEV assay could represent a solution to existing challenges in functional assays for EVs.

## Materials and Methods

Nanoalgosomes and cell derived-EVs are isolated and characterized as previously described in Adamo G. et al., 2021, and the relative methods are reported in Supporting methods.

### Set up of functional enzymatic assay: pilot test

Fluorescein diacetate (FDA, Sigma-Aldrich) stock solution was prepared in acetone at a final concentration of 2 mg/mL. 2 x 10^10^ nanoalgosomes were incubated with 35 µmol of fluorescein diacetate, reaching a final volume of 200 µl with 0.2 µm filtered PBS without Ca^++^ and Mg^++^. The same amount of fluorescein diacetate was added to 0.2 µm filtered PBS without Ca^++^ and Mg^++^ (nanoalgosome-vehicle). Fluorescence emission was measured every 15 minutes, up to 16 h, using a blue filter of the GloMax Discover Microplate Reader.

### Preparation fluorescein standards curve

Fluorescein sodium salt (ex/em 490/514 nm, Sigma-Aldrich) was used to generate a standard calibration curve.

The fluorescence intensity of serial dilutions of fluorescein (from 300 nmol to 0 nmol), diluted in PBS without Ca^++^ and Mg^++^ at a final volume of 200 µl, was measured in a 96-well-plate using a blue filter of the GloMax Discover Microplate Reader (Promega).

### Measure of enzymatic activity

To calculate the specific enzymatic activity, the enzyme units (U), the amount of esterases that catalyses the reaction of 1 nmol of substrate into product per minute, was determined. EV enzymatic activity values were expressed in U as nanomoles of fluorescein produced per minute (nmol/min). This value was determined by fitting the fluorescence intensities measured at the end of the functional assay to the fluorescein concentration of the standard curve. Subsequently, this value was divided by the total assay time (180 minutes) to obtain the nanomoles per minute.

### Michaelis–Menten Kinetic

2 x 10^10^ nanoalgosmes were incubated with different concentrations of fluorescein diacetate (0, 1.6, 3.3, 6.6, 13, 26 x 10^3^ nmol) dissolved in 0.2 µm filtered PBS without Ca^++^ and Mg^++^, in a final volume equal to 200 µl. The fluorescence emission was followed until 3 hours, in a 96-well-plate using a blue filter of the GloMax Discover Microplate Reader (Promega). Obtained values were corrected by subtracting background fluorescence value (i.e., PBS with fluorescein diacetate). Fitting these data to the standard calibration curve, the Michaelis–Menten equation was applied using nonlinear regression and plotting the reaction’s initial rates (nmol/min) as a function of fluorescein diacetate concentration. Vmax and Km were found using GraphPad software.

### Analytical sensitivity

Serial dilutions of nanoalgosomes (from 2 x 10^10^ to 3 × 10^8^ sEVs) were incubated with 18 µmol of fluorescein diacetate, reaching a final volume of 200 µl with 0.2 µm filtered PBS without Ca^++^ and Mg^++^. Fluorescence emission was followed up to 3h, using a blue filter of the GloMax Discover Microplate Reader. The same amount of fluorescein diacetate was added to 0.2 µm-filtered PBS without Ca^++^ and Mg^++^ (nanoalgosome-vehicle). This background fluorescence value was subtracted from the fluorescence intensity values of each sample, for all the condition described. The relative methods of fluorescein diacetate auto-hydrolysis in different media are reported in Supporting methods.

### Analytical specificity

2x10^10^ nanoalgosomes were incubated with 0.1, 0.25, 0.5 and 1% Triton-X100 (Sigma-Aldrich) for 30 min at room temperature. The same amount of nanoalgosomes were boiled at 100°C for 10 min. Next, samples were incubated with 18 µmol of fluorescein diacetate, reaching a final volume of 200 µl with 0.2 µm filtered PBS without Ca^++^ and Mg^++^. Fluorescence emission was followed up to 3h, and enzymatic activity was determined as described previously. As control, the same amount of fluorescein diacetate was added to 0.2 µm-filtered PBS without Ca^++^ and Mg^++^ boiled at 100°C for 10 min or with 0.1, 0.25, 0.5 and 1% Triton-X 100. This background fluorescence value was subtracted from the fluorescence intensity values of each respective sample.

Additionally, 2x10^10^ nanoalgosomes were incubated with 18 µmol of fluorescein diacetate, reaching a final volume of 200 µl with 0.2 µm filtered PBS without Ca^++^ and Mg^++^. During the time course of the detectEV assay, specifically after 1, 2, and 3 hours post fluorescein diacetate addition, nanoalgosomes were treated with 1% Triton-X 100. Fluorescence emission was measured every 15 minutes, up to 195 minutes, using a blue filter of the GloMax Discover Microplate Reader. As control, the same amount of fluorescein diacetate was added to 0.2 µm-filtered PBS with/without Ca^++^ and Mg^++^ with 1% Triton X-100. This background fluorescence value was subtracted from the fluorescence intensity values of each respective sample.

For protease treatment, 10^11^ nanoalgosomes were incubated with a 0.25% (w/v) trypsin solution (Invitrogen, Life Technologies) at 37 °C for 15 minutes. An EV-free control was prepared by using 0.2 µm filtered PBS without Ca++ and Mg++. After the digestion period, both EV samples and the negative control underwent a washing step to remove trypsin. Given that FDA can be hydrolyzed by proteases, it was essential to eliminate trypsin post-treatment. This was achieved through five washes with PBS using a 100 kDa Amicon filter at low speed (3000 g for 5 minutes, repeated 5 times at 4°C). The absence of trypsin activity was confirmed in the negative control before proceeding with the detectEV assay on the protease-treated EV-samples.

### Comparative Analysis of Esterase-Sensitive Probes

For the comparative study, we used esterase-sensitive probes, including carboxyfluorescein diacetate succinimidyl ester (CFSE), calcein acetoxymethyl ester (calcein-AM) (both from Sigma-Aldrich), and the lipophilic dye PKH67 (Sigma-Aldrich) as an esterase-sensitive negative control. Like fluorescein diacetate, calcein-AM is hydrolyzed by esterases to produce a polar green-fluorescent product, while CFSE is converted by esterases into an intermediate that interacts with amines to generate a highly fluorescent green dye. PKH67, a green lipophilic dye, integrates into the membrane without requiring enzymatic activation. Specifically, 2×10^10^ nanoalgosomes were incubated with 10 µM calcein-AM, 10 µM CFSE (CFDA-SE), and 10 µM PKH67, respectively. EV-free PBS solution, used as a negative control, was incubated with the same probes to monitor background signal.

### Fluorescence-based microplate reader analysis

For the microplate reader analysis, 2×10^10^ nanoalgosomes were incubated with each probe, with and without detergent pre-treatment (using 1% Triton X-100 as described before), reaching a final volume of 200 µL with 0.2 µm filtered PBS without Ca^++^ and Mg^++^. The same amount of each probe was added at the same concentration to 0.2 µm filtered PBS without Ca^++^ and Mg^++^ (used as negative control that correspond to EV-free PBS) with and without 1% Triton X-100 . Fluorescence emission was measured every 15 minutes, up to 3 hours, using a blue filter 488 nm on the GloMax Discover Microplate Reader. The background fluorescence value of the negative control was subtracted from the fluorescence intensity values of each respective sample.

### Flow Cytometry

All flow cytometry experiments were performed using a CytoFLEX SRT Cell Sorter (Beckman Coulter), operated with CyExpert SRT software. The CytoFLEX SRT was initialized according to the manufacturer’s recommendations to ensure optimal performance. The CytoFLEX SRT used in this study was equipped with three lasers: a red laser (638 nm), a blue laser (488 nm), and a violet laser (405 nm). For the analysis of small particles such as EVs, the configuration was adjusted for violet side scatter (VSSC) detection using the violet (405 nm) laser, and fluorescence detection was performed with the blue laser (488 nm). Megamix-Plus FSC beads (BioCytex), consisting of distinct populations with sizes of 100, 160, 200, 240, 300, 500, and 900 nm, were used to define the size range for EV detection. These beads helped establish the gating strategy for accurately identifying and analyzing EVs based on their size. EV samples were diluted 1:20 in 0.2 µm filtered PBS without Ca^++^ and Mg^++^ to a final volume of 1 mL. To create a stable and slow-velocity core stream, which is recommended for the detection of EVs, the sample acquisition speed was adjusted. The flow rate was set to 10 µL per minute, ensuring precise and consistent sample introduction into the flow cytometer. For each sample, at least 10,000 events were recorded. The acquisition rate was maintained at approximately 6,000 events per second to ensure a high-resolution analysis.

### detectEV assay for human cell-derived sEVs

2x10^10^ sEVs isolated from HEK 293T cells were employed to conduct a time course measurements. After the incubation with 18 µmol of fluorescein diacetate, the fluorescence emission was measured every 15 minutes, up to 180 minutes, using a blue filter of the GloMax Discover Microplate Reader. As negative control, the same amount of fluorescein diacetate was added to 0.2 µm-filtered PBS without Ca^++^ and Mg^++^. Next, serial dilutions of HEK 293T derived sEVs (from 2 x 10^10^ to 3 × 10^8^ sEVs) were incubated with 18 µmol of fluorescein diacetate, reaching a final volume of 200 µl with 0.2 µm filtered PBS without Ca^++^ and Mg^++^. Fluorescence emission was followed up to 3h, using a blue filter of the GloMax Discover Microplate Reader. The same amount of fluorescein diacetate was added to 0.2 µm-filtered PBS without Ca^++^ and Mg^++^ (sEV-vehicle). This background fluorescence value was subtracted from the fluorescence intensity values of each sample, for all the condition described. Further, 2x10^10^ HEK 293T derived sEVs were incubated with 1% Triton-X 100 for 30 min at room temperature. Next, EV-samples were incubated with 18 µmol of fluorescein diacetate, reaching a final volume of 200 µl with 0.2 µm filtered PBS without Ca^++^ and Mg^++^. Fluorescence emission was followed up to 3h, and enzymatic activity was determined as described previously. As control, the same amount of fluorescein diacetate was added to 0.2 µm-filtered PBS without Ca^++^ and Mg^++^ with 1% Triton-X 100. This background fluorescence value was subtracted from the fluorescence intensity values of each respective sample. For MDA-MB231 and 1-7 HB2 cell derived EVs, 2x10^10^ sEVs were employed to conduct the detectEV assay, using the same experimental setting described before.

### Application of detectEV assay

#### Isolation methods

2x10^10^ nanoalgosomes of the same *T. chuii* conditioned media, isolated by dUC and TFF, were used to perform the functional enzymatic assay, using the same settings described before (Adamo et al., 2021). A detailed description of methods used for EV isolation are reported in Supporting methods.

#### Storage conditions

Nanoalgosomes were stored for 10 days at different condition: 4°C, -20°C, −80 °C and upon lyophilization in 5% and 8% sucrose, at 4°C for a week. Lyophilized samples were carefully rehydrated in 0.2 µm-filtered Milli-Q water. After NTA analysis on all stored samples, detectEV assay was performed, as described previously.

#### Loading methods

2x10^12^ nanoalgosomes (and nanoalgosome-vehicle*, i.e*. 0.2 µm filtered PBS without Ca^++^ and Mg^++^, used as a negative control) underwent the subsequent treatments: -Electroporation was performed in Gene Pulser cuvettes (0.4 cm cell electrode gap) on a BioRad Gene Pulser equipped with a capacitance extender, with two conditions selected, E1 (125 μF, 400 V and 2 pulse time of 20 ms) and E2 (125 μF, 250 V and 2 pulse time of 30 ms). **-**Sonication was performed using two settings: ultrasonic probe sonicator (S1) with 20% amplitude for six cycles of 30 seconds on/off for a total of 3 min, with 2 min cooling, then incubation for 60 min at 37°C; ultrasonic bath (S2) 40 KHz, 40% amplitude for 2 cycles of 30 seconds on/off, then incubation for 60 min at 37°C.

-Freeze–Thaw was performed in 3 cycles of freezing at -80° for 30 min and thawing at room temperature (RT) for 30 min.

-Saponin treatment was performed using two settings: incubation with 0.1 mg/mL (Sap1) and 0.002 mg/mL (Sap2) of saponin (Sigma-Aldrich) at RT for 10 min.

-Extrusion using a mini-extruder equipped with polycarbonate membrane filters of 100 nm and 200 nm pore size (Avestin, Manheim, Germany). Each sample was extruded 31 times.

After the application of each method, NTA analysis was performed on all samples. Next, the enzymatic functional assay was performed, as described previously.

#### Batch-to-batch reproducibility

2 x 10^10^ nanoalgosomes from different batches (named Alg1, Alg2, Alg3 and Alg4) were incubated with 18 µmol of fluorescein diacetate, reaching a final volume of 200 µl with 0.2 µm filtered PBS without Ca^++^ and Mg^++^. Fluorescence emission was followed up to 3h, and enzymatic activity was determined as described above. As negative controls, nanoalgosomes were boiled at 100°C for 10 min.

#### Antioxidant activity assay

Intracellular ROS levels in living cells were assessed by employing 2′, 7′-dichlorofluorescein diacetate (DCF-DA; Sigma-Aldrich). DCF-DA undergoes oxidation to form fluorescent DCF (2′, 7′-dichlorofluorescein) in the presence of reactive oxygen species (ROS), enabling detection using a spectrofluorometer. The antioxidant assay was conducted on 1-7 HB2 cell line. Specifically, 4x10^3^ cells were plated in 96-well microplates for 24 hours. Subsequently, cells were treated with Alg1, Alg3 and Alg4 (0.5 μg/mL, 10^10^ EVs/mL) for an additional 24 hours. Following removal of the medium, cells were exposed to PBS containing 40 μM of DCF-DA and incubated in a humidified atmosphere (5% CO2 at 37°C) for 1 hour. The cells were then subjected to treatment with or without the oxidative agents TBH (250 μM for 1 hour) (tert-butyl hydroperoxide solution, Sigma-Aldrich), in the absence or presence of nanoalgosomes. Untreated cells served as a control to establish the baseline intracellular ROS percentage. After thorough washing steps, fluorescence intensity was measured using a fluorescence plate reader with an excitation wavelength of 485 nm and an emission wavelength of 538 nm (GloMax® Discover Microplate Reader, Promega). The relative percentage of intracellular ROS was normalized respect to untreated cells (control).

### Statistical analysis

GraphPad Prism version 10 was used for statistical analysis. T-Test, One-way analysis of variance (ANOVA) or Two-way analysis of variance (ANOVA) followed by Tukey’s multiple comparisons were performed when two, three or more than three means of independent groups were compared, respectively. Statistical significance was set as p < 0.05 and star significance were distributed as * for p < 0.05, ** for p < 0.01, *** for p < 0.001, and **** for p < 0.0001, while ns correspond to non-significant differences. Each measurement reported here was repeated as experimental triplicate, and mean values, as well as standard deviations, were calculated.

## Author Contributions (CRediT)

**Giorgia Adamo:** Conceptualization, Investigation, Methodology, Formal analysis, Visualization, Validation, Funding acquisition, Writing - Original Draft and Final Review & Editing; **Sabrina Picciotto:** Methodology, Formal analysis, Data curation, Software, Writing - Original Draft; **Paola Gargano:** Formal analysis, Data curation, Software, Writing - Original Draft; **Angela Paterna**: Formal analysis, Data curation; **Samuele Raccosta**: Formal analysis, Data curation; **Estella Rao**: Formal analysis, Data curation; **Daniele Paolo Romancino**: Formal analysis, Data curation, **Giulio Ghersi:** Data curation, Final Review; **Mauro Manno**: Formal analysis, Data curation; Final Review; **Monica Salamone:** Formal analysis, Data curation; **Antonella Bongiovanni**: Conceptualization, Investigation, Validation, Writing - Original Draft and Review & Editing, Supervision, Visualization, Project administration, Funding acquisition; Final Review & Editing.

## Associated Content

Supporting information (PDF) contains additional figures and detailed supporting methods; this file is available free of charge.

Supplemental material online includes additional figures related to immunoblot analysis for EV biomarkers, available via Figshare (https://figshare.com/s/33f82148a7f774446c63).

## Supporting Methods

### Microalgae cultivation

A stock culture of the microalgae *T. chuii* (CCAP 66/21b), grown in borosilicate glass bottles was used to start new cultures in bottles via a 25% v/v inoculum. Microalgae medium are constitute of certified ultrafiltered, clear and sterile marine water (Steralmar) with Guillard′s (F/2) enrichment solution (F/2 medium). Cultures were kept for 4 weeks at a temperature of 22°C ± 2°C under continuous air flow and exposed to white light with a photoperiod of 14 h light and 10 h dark. Bottles were gently shaken every 2 days in order to homogenize the cultures. Microalgae were cultured in sterile conditions by using 0.22 μm filters at the bottle inlets. Cell growth was monitored every week by optical density at 600 nm and cell counting.

### Mammalian cell cultivation

The following cell lines were used for EV production: (i) 1–7 HB2, normal mammary epithelial cells; (ii) MDA-MB 231, human breast cancer epithelial cells; and (iii) HEK 293T, human embryonic kidney cells. MDA-MB-231 and HEK293T were cultured in Dulbecco’s Modified Eagle Medium (DMEM, Sigma-Aldrich) containing 10% (v/v) Fetal Bovine Serum (FBS, Gibco, Life Technologies) plus 2 mM L-glutamine, 100 U/ml Penicillin and 100 mg/ml Streptomycin (Sigma-Aldrich); instead, the 1–7 HB2 cell line was cultured in low-glucose DMEM (Sigma-Aldrich) containing 10% (v/v) Fetal Bovine Serum, plus 2 mM L-glutamine, 100 U/ml Penicillin and 100 mg/ml Streptomycin, 5 μg/ml Hydrocortisone (Sigma-Aldrich) and 10 μg/ml Bovine Insulin (Sigma-Aldrich). All cell lines were maintained at 37°C in a humidified atmosphere (5% CO_2_). All cell lines were seeded in 175 cm² flasks until 70% confluence was reached. Next, cells were washed twice with phosphate saline buffer (PBS, Sigma-Aldrich) with Ca^++^ and Mg^++^ and cultured in appropriate medium supplemented with 10% of EV-depleted FBS. After 48 h, sEVs were isolated from each cell-conditioned medium.

### Microalgae and human-derived-EV isolation

#### Differential ultracentrifugation

The isolation of both human cell and microalgal derived-EVs was performed by dUC, as described in Picciotto et al., 2021. Briefly, conditioned media was subjected to serial centrifugation steps at 4 °C. The first was at 300 x g for 10 min. The collected supernatant was subjected to 2,000 x g for 10 min to remove all cell debris. Large EVs (lEVs) were separated in 50 ml eppendorf polypropylene conical tubes at 10,000 × g for 30 min at 4°C using an eppendorf rotor F34-6-38. We then collected sEVs from the supernatant into Beckman Coulter polypropylene open top tubes via centrifugation at 118,000 × g for 70 min at 4°C using a Beckman SW28 rotor. After a PBS washing step, the pellet was re-suspended in PBS without Ca^++^ and Mg^++^ for subsequent analyses.

#### Tangential flow filtration

Isolation of microalgal derived-EVs was performed using the KrosFlo® KR2i TFF System from Repligen (Spectrum Labs., Los Angeles, CA, USA) and three modified polyethersulfone hollow fiber membranes (S04-E65U-07-N, S04-P20-10-N and S04-E500-10-N, Spectrum Labs). Briefly, microalgae cultures were purified by microfiltration and ultra-filtration using a hollow fiber cartridge housed in the KrosFlo® KR2i, with cut-offs of 650 nm (S04-E65U-07-N, Spectrum Labs), 200 nm (S04-P20-10-N, Spectrum Labs), and 500-kDa (S04-E500-10-N, Spectrum Labs). Feed flow and transmembrane pressure (TMP) were kept constant at 450 ml/min and 0.05 bar, respectively. The small and large EVs recovered from the retentate of the 500 kDa and 200 nm cut-off TFF filter modules, respectively, were concentrated until a final volume of approximately 150 ml. Subsequently, using a smaller 500 kDa cut-off TFF filter module (C02-E500-10-N, Spectrum Labs, MicroKros) with a feed flow of 75 ml/min and a permeate flow of 2 ml/min, samples were further concentrated and diafiltrated seven times with PBS, without Ca^++^ and Mg^++^, reaching a final volume of approximately 5 ml.

### EV characterization

#### Nanoparticle tracking analysis (NTA)

Particle concentration and size distributions of EVs were measured using an NTA instrument (NanoSight NS300, Malvern Panalytical, UK). EV samples were diluted to generate a dilution in which 20–120 particles per frame were tracked, to obtain a concentration within the recommended measurement range (1 - 10 × 10^8^ particles/ml). For each individual sample, five videos of 60 s were recorded and analyzed with detection threshold 5 and camera level 15-16. All videos were analyzed by NTA Software.

#### Dynamic light scattering (DLS)

An aliquot of the vesicle solution was pipetted and centrifuged at 1000×g for 10 minutes at 4 °C to eliminate any dust particles. The supernatant was withdrawn using clean pipet tips, placed directly into a quartz cuvette, and incubated at 20 °C in a thermostated cell compartment of a Brookhaven BI200-SM goniometer.. The g_2_(t) autocorrelation function of scattered light intensity was measured at a scattering angle ϑ = 90° using a Brookhaven BI-9000 correlator. g_2_(t) is related to the size D_h_ of diffusing particles and their size distribution P(D_h_). The relation is given by g_2_(t) = 1 + |β ∫ P(D_h_) exp[−D(D_h_)q^2^ t]|^2^, where β is an instrumental parameter, q = 4πñλ^-1^ sin[ϑ/2] is the scattering vector, with ñ being the refractive index of the medium (ñ=1.3336), and D(D_h_) is the diffusion coefficient of a particle with a hydrodynamic diameter D_h_, determined by the Stokes-Einstein relation D(D_h_) = k_B_T [3πηD_h_]⁻¹, with T being the temperature and η the medium viscosity. The size distribution P(D_h_) is calculated assuming that the diffusion coefficient distribution is shaped as a Schultz distribution.Two robust parameters derived from this analysis are the z-averaged hydrodynamic diameter D_z_ and the polydispersity index PDI, an estimate of the distribution width.

#### Protein quantification and Immunoblot analysis

Microalgae lysates were prepared as described in Adamo et al, 2021; mammalian cellular lysates were prepared by RIPA buffer extraction protocol. RIPA lysis buffer was prepared based with 25 mM Tris-HCl pH 7.6, 150 mM NaCl, 1% NP-40, 1% sodium deoxycholate, 0.1% SDS, enriched with EDTA to a final concentration of 1 mM, a protease inhibitor cocktail, and a phosphatase inhibitor cocktail. After 15 minutes, the protein lysates were centrifuged at 10,000× g, for 20 min at 4 °C and the protein-rich supernatants stored at −80 °C until further processing Total protein of EV and cellular lysates was quantified using the microBCA protein assay kit (Thermo Fisher Scientific) according to the manufacturer’s instructions, using bovine serum albumin (BSA) as a standard. Then, EV and lysate samples and standards were mixed with the working reagent and incubated for 1 h at 37°C. Absorbance values were measured at 562 nm, using a GloMax® Discover Microplate Reader. Next, EVs and cell lysates were separated by sodium dodecyl-sulfate polyacrylamide gel electrophoresis (SDS-PAGE) (10%). A total of 10 μg of EVs samples (in PBS) were mixed with proper volumes of 5X loading buffer (0.25 M Tris-Cl pH 6.8, 10% SDS, 50% glycerol, 0.25 M dithiothreitol (DTT), 0.25% bromophenol blue). Then, the samples were heated at 100°C for 5 minutes and loaded onto a 10% sodium dodecyl sulfate-polyacrylamide gel for electrophoretic analyses. Proteins were blotted onto polyvinylidene fluoride membranes (PVDF), which were blocked with 3% bovine serum albumin (BSA) in TBS-T solution (50 mM Tris HCl pH 8.0, 150 mM NaCl with 0.05% Tween 20) for 1 hour at room temperature. For nanoalgosome immunoblot were used the following primary antibodies: anti-H+-ATPase (dil. 1:1000 in 3% BSA/TBS-T1x, Clonality: Polyclonal, Agrisera), anti-Alix (clone 3A9, dil. 1:150 in 3% BSA/TBS-T 1X, sc-53538. Santa Cruz), anti-ß-actin (clone AO480, dil. 1:500 in 3% BSA/TBS-T 1X, Millipore); for mammalian cell derived-EV immunoblot were used the following primary antibodies: anti-Enolase (clone A5, dil. 1:400 in 3% BSA/TBS-T1X, sc-271384. Santa Cruz), anti-CD63 tetraspanin (polyclonal, dil. 1:500 in 3% BSA/TBS-T 1X, Invitrogen), anti-HSP70 (W27,sc-24, dil. 1:150 in 3% BSA/TBS-T 1X, Santa Cruz) and anti-β-actin (AC-15, dil. 1:150 in 3% BSA/TBS-T 1X, Santa Cruz). Each primary antibody was incubated for 1 hour at room temperature. After washing, the membrane was incubated for 1 hour with secondary antibodies according to the manufacturer’s instructions (horseradish peroxidase-conjugated secondary anti-mouse or anti-rabbit antibodies, Cell Signaling). The membrane was washed four times in TBS-T for 20 minutes. The immunoblots were revealed using SuperSignal Pierce ECL (Thermo Fisher Scientific).

#### Atomic Force Microscopy (AFM)

Sample preparation: glass slides were cleaned by boiling acetone, dried by nitrogen stream and exposed to UV rays (30 W Hg lamp); then they have been i) treated with a 0.25 M (3-aminopropyl)-triethoxysilane in chloroform solution for 3 minutes, and then rinsed with chloroform and dried with nitrogen; (ii) treated with 0.4 M glutaraldehyde aqueous solution for 3 minutes and then rinsed with Milli-Q water and dried with nitrogen. The vesicle solutions diluted in PBS to a final concentration of a few μg/ml were deposited onto APTES/glutaraldehyde functionalized glass slides and incubated overnight. Then, the samples were gently rinsed by PBS to remove non-adsorbed vesicles.

Vesicle imaging: Quantitative Imaging AFM measurements were carried out in PBS by using a Nanowizard III scanning probe microscope (JPK Instruments AG, Germany) equipped with a 15-μm z-range scanner, and AC40 (Bruker) silicon cantilevers (spring constant 0.1 N/m, typical tip radius 8 nm). The images (resolution 206x206 pixels) were acquired at force setpoint 110 pN and extend speed 25 μm/s (z-length 50 nm or 80 nm and pixel time 5 ms or 8 ms respectively for algosomes and EVs from mammalian cells). The cantilever was thermally calibrated by using the tool in JPK software (Hutter and Bechhoefer, 1993).

#### Fluorescent-nanoparticle tracking analysis (F-NTA)

EVs were stained with 500 nm of 4-(2-[6-(diethylamino)-2-naphthalenyl]ethynyl)-1-(3 sulfopropyl) pyridinium, DI-8-ANEPPS (Ex/Em: 467/631 nm, Thermo Fisher Scientific), previously filtered by 20 nm filters (Whatman Anotop filters), whose fluorescence is activated in a-polar environments, and specifically enhanced when bound to the lipid membrane of EVs, with a higher quantum yield concerning any binding to hydrophobic protein regions.

This makes the Di-8-ANEPPS fluorescent signal highly EV-specific. After 1 h at room temperature, F-NTA analyses were carried out as described in Adamo et al., 2021.

### Fluorescein diacetate auto-hydrolysis background analysis

To assess potential FDA auto-hydrolysis, background fluorescence measurements were performed in clear and sterile marine water (Steralmar) with Guillard′s (F/2) enrichment solution (F/2 medium) and Dulbecco’s Modified Eagle Medium (DMEM, Sigma-Aldrich) containing 10% (v/v) EV-depleted Fetal Bovine Serum (FBS, Gibco, Life Technologies) plus 2 mM L-glutamine, 100 U/ml Penicillin and 100 mg/ml Streptomycin (Sigma-Aldrich); instead, low-glucose DMEM (Sigma-Aldrich) containing 10% (v/v) EV-depleted Fetal Bovine Serum, plus 2 mM L-glutamine, 100 U/ml Penicillin and 100 mg/ml Streptomycin, 5 μg/ml Hydrocortisone (Sigma-Aldrich) and 10 μg/ml Bovine Insulin (Sigma-Aldrich). Fluorescein diacetate auto-hydrolysis background was monitored in each media both in their original formulation and in EV-free post-purification forms. Specifically, F/2 medium was purified using the TFF, while the DMEM media using dUC, and resuspended in 0.2µm filtered PBS. Then, detectEV assay was applied in each samples, using the same experimental setting described for EVs.

---

**Supporting Figure 1.**
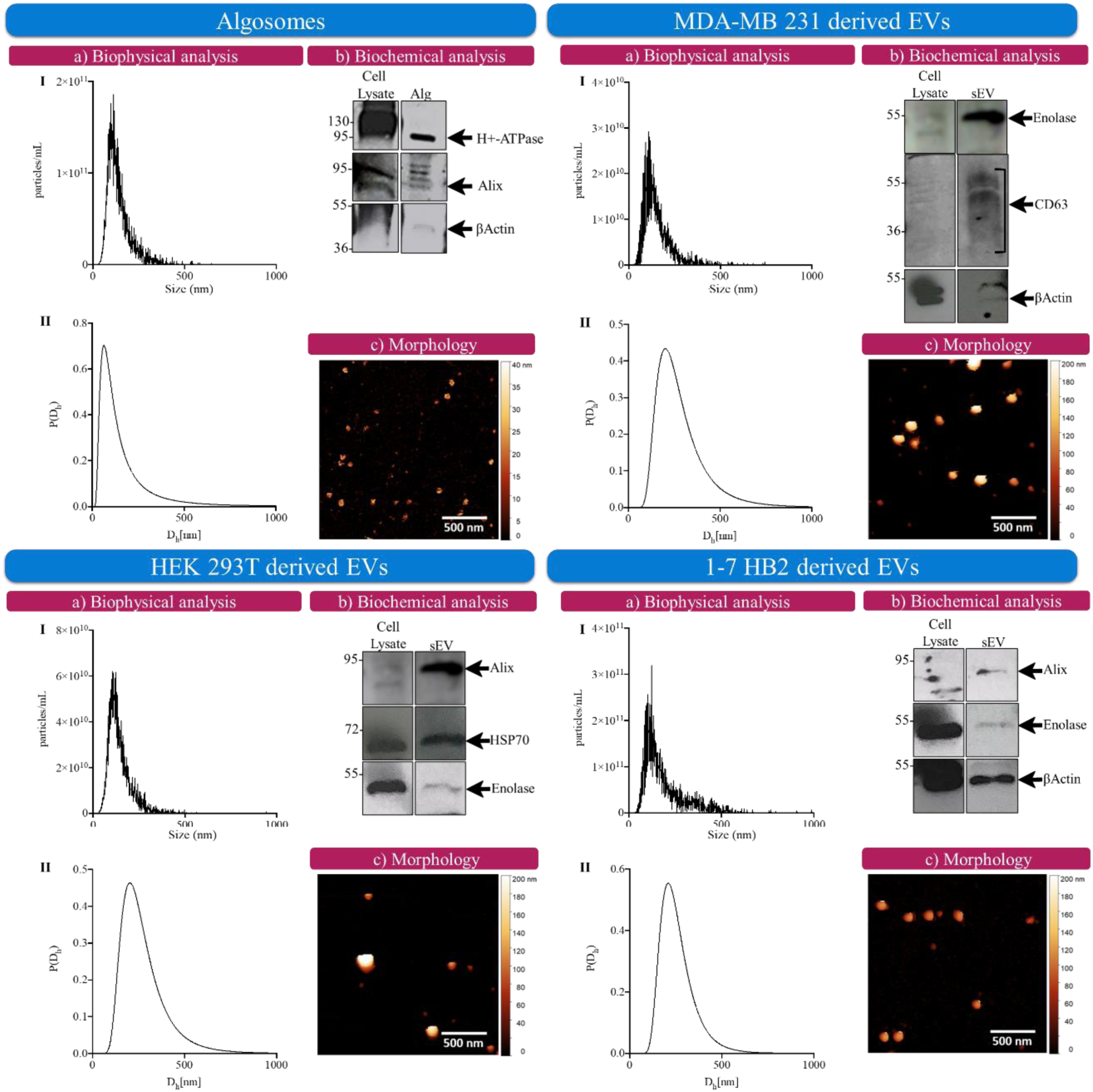
Characterization of nanoalgosomes and sEVs derived from MDA-MB 231, HEK 293T, and 1-7 HB2 cells. a) Biophysical analyses included (I) representative NTA measurements and (II) Dynamic Light Scattering (DLS) showing the particle size distribution of the vesicles (error bars represent standard deviations obtained from five measurements of the same sample); b) Biochemical analyses: immunoblotting for specific EV biomarkers performed on sEVs and the corresponding cell lysates. The biomarkers include common EV markers to confirm the presence and purity of EVs in the samples. c) Morphological analyses: representative Atomic Force Microscopy (AFM) images depicting the morphology of the vesicles. The scale bar indicates 500 nm for all images, providing a visual representation of the vesicle size and structure.

**Supporting Figure 2.**
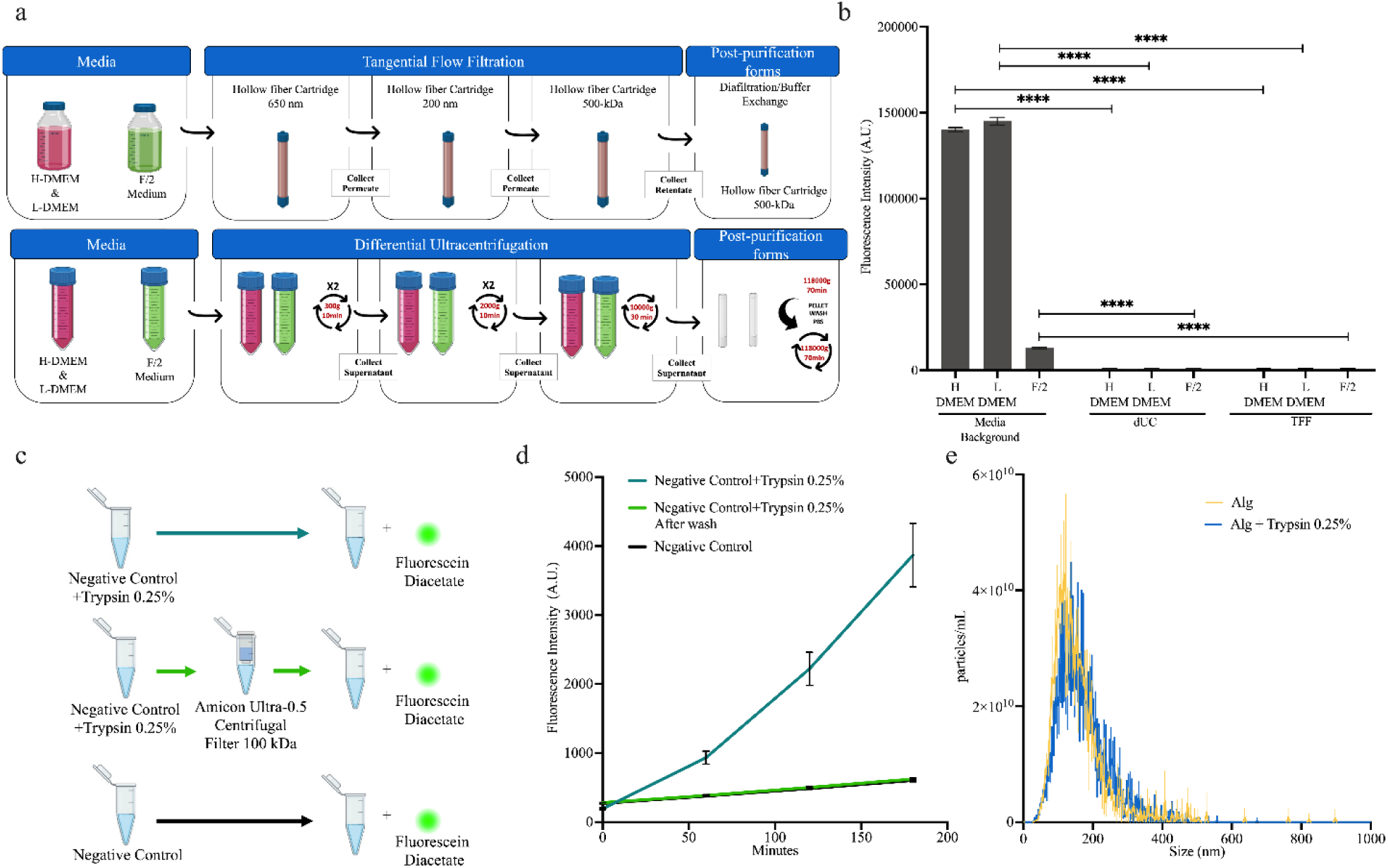
a-b) Fluorescein diacetate auto-hydrolysis background analysis in EV-free media. a) Schematic of the experimental setup: fluorescein diacetate auto-hydrolysis background was monitored in *Tetraselmis chuii* medium (*e.g.*, F/2) and in MDA-MB 231, HEK 293T, and 1-7 HB2 cell media (*e.g.*, complete high and low-glucose DMEM, H- and L-DMEM) in their original formulation (*e.g.*, media) and in their dUC- and TFF-based post-purification forms. Specifically, media were purified using dUC and TFF and resuspended in 0.2µm filtered PBS. b) detectEV assay was applied to each samples, using the same experimental setting described for EVs. Error bars represent the standard deviation of the mean of independent experiments (n= 3). Two-way ANOVA was used to assess the statistical significance of differences between media background and their dUC- and TFF-based post-purification forms showing ****p<0.0001. **c-e) Experimental Validation of Trypsin Removal and Substrate Hydrolysis.** c) Schematic of the experimental setup to validate the effectiveness of 100 kDa Amicon filter washing in removing 0.25% trypsin after the digestion of EV-samples. In this setup, PBS (the negative control) was incubated with trypsin for 15 minutes, followed by fluorescein diacetate. The tested conditions included also the negative control with trypsin, followed by washing steps and incubation with fluorescein diacetate, and the negative control incubated with fluorescein diacetate only. d) Fluorescence intensity was measured by plate reader over time (up to 180 minutes) for the samples described in (c). The fluorescence data demonstrate that the washing step effectively removes trypsin, as indicated by the reduced fluorescence in the washed samples compared to the unwashed ones in which the increase in fluorescence intensity indicates the hydrolysis of fluorescein diacetate by trypsin (n=3 independent experimental replicates). e) Nanoparticle Tracking Analysis of nanoalgosomes after protease treatment: NTA measurements were conducted to assess the size distribution and concentration of nanoalgosomes before and after a 15 minutes digestion with 0.25% trypsin at 37°C. The results revealed no significant differences in size distribution and concentration between the treated and untreated samples, indicating that the trypsin digestion does not alter this physical property of the nanoalgosomes. The analysis is based on five independent measurements for each samples.

**Supporting Figure 3.**
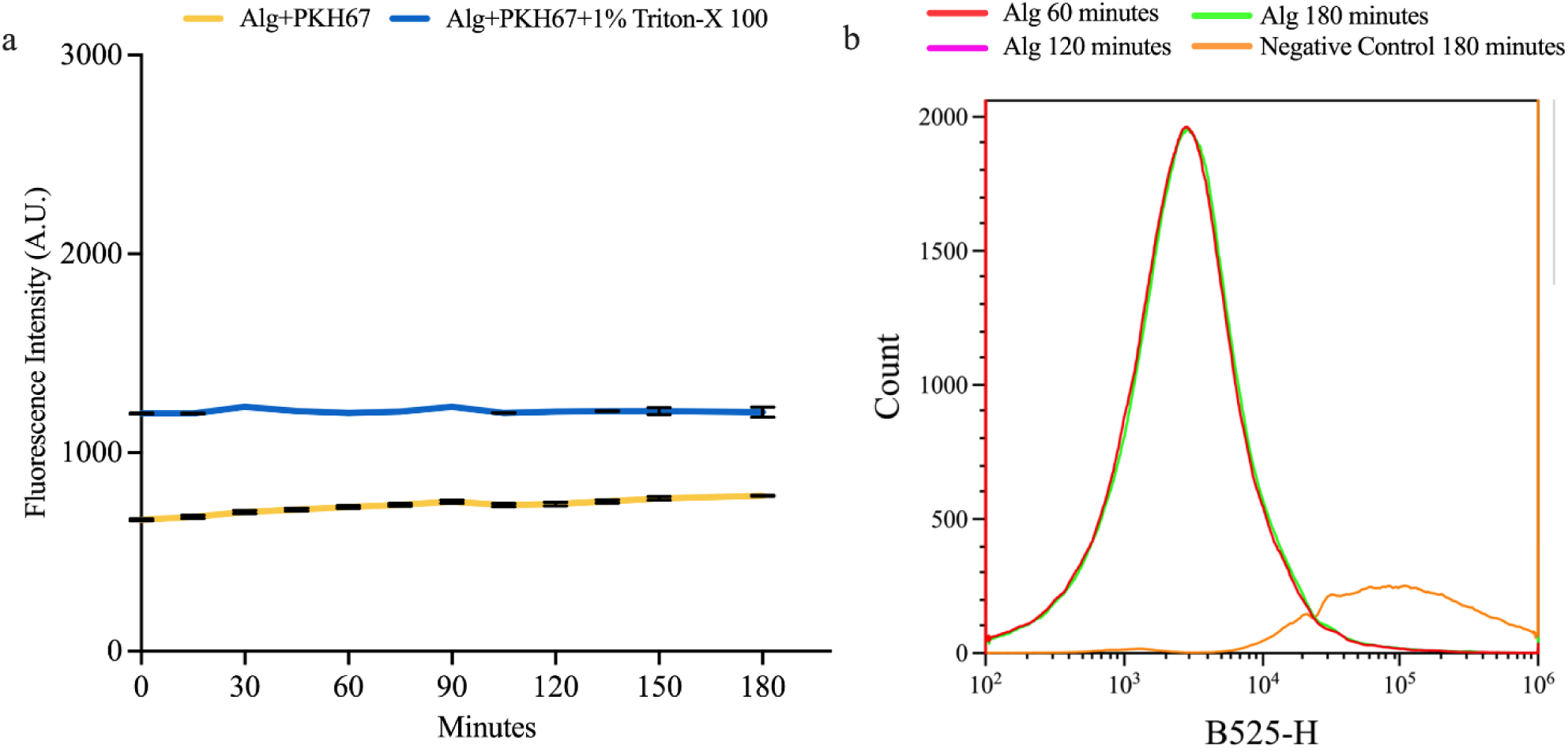
Analysis of PKH67 fluorescence intensity measured by (a) fluorescence-based plate reader and (b) flow cytometry. a) The graph show the fluorescence intensity expressed in arbitrary unit, A.U., (excitation 488nm, emission 520nm) up to 180 minutes for nanoalgosomes treated or not with 1% Triton-X 100 and incubated with PKH67. Error bars represent the standard deviation of the mean of the fluorescence intensity subtracted from the respective background signals measured in the corresponding negative controls, which is PBS with PKH67, with and without 1% Triton-X 100. Three independent experiments were considered (n=3). b) Flow cytometry analyses: histograms of counts versus fluorescence (blue laser 488 nm) of nanoalgosome incubated 60, 120, and 180 minutes with PKH67. Negative control samples correspond to PBS incubated 180 minutes with PKH67.

**Supporting Figure 4.**
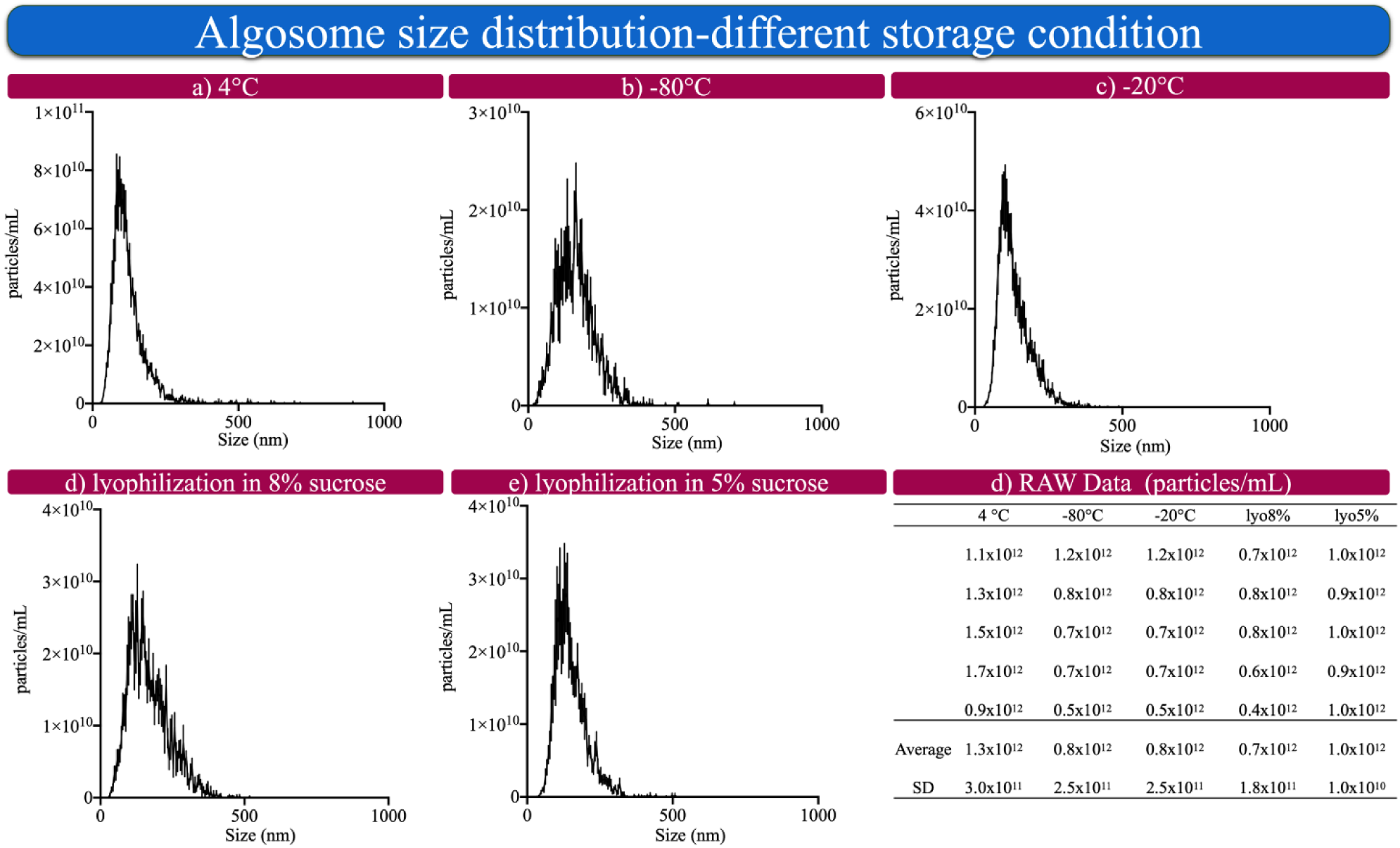
Nanoalgosome size distribution by NTA under five distinct storage conditions. a) 4°C, b) -80°C, c) -20°C, d) lyophilization in 8% sucrose, and e) lyophilization in 5% sucrose. Each histogram represents the average of five NTA measurements and error bars represent standard deviations. f) Table presents the raw particle count data (particles/mL) for nanoalgosomes stored under different conditions. Each condition was measured in five replicates, with the particle counts provided for each replicate. The average particle count and standard deviation (SD) for each storage condition are also included.

**Supporting Figure 5.**
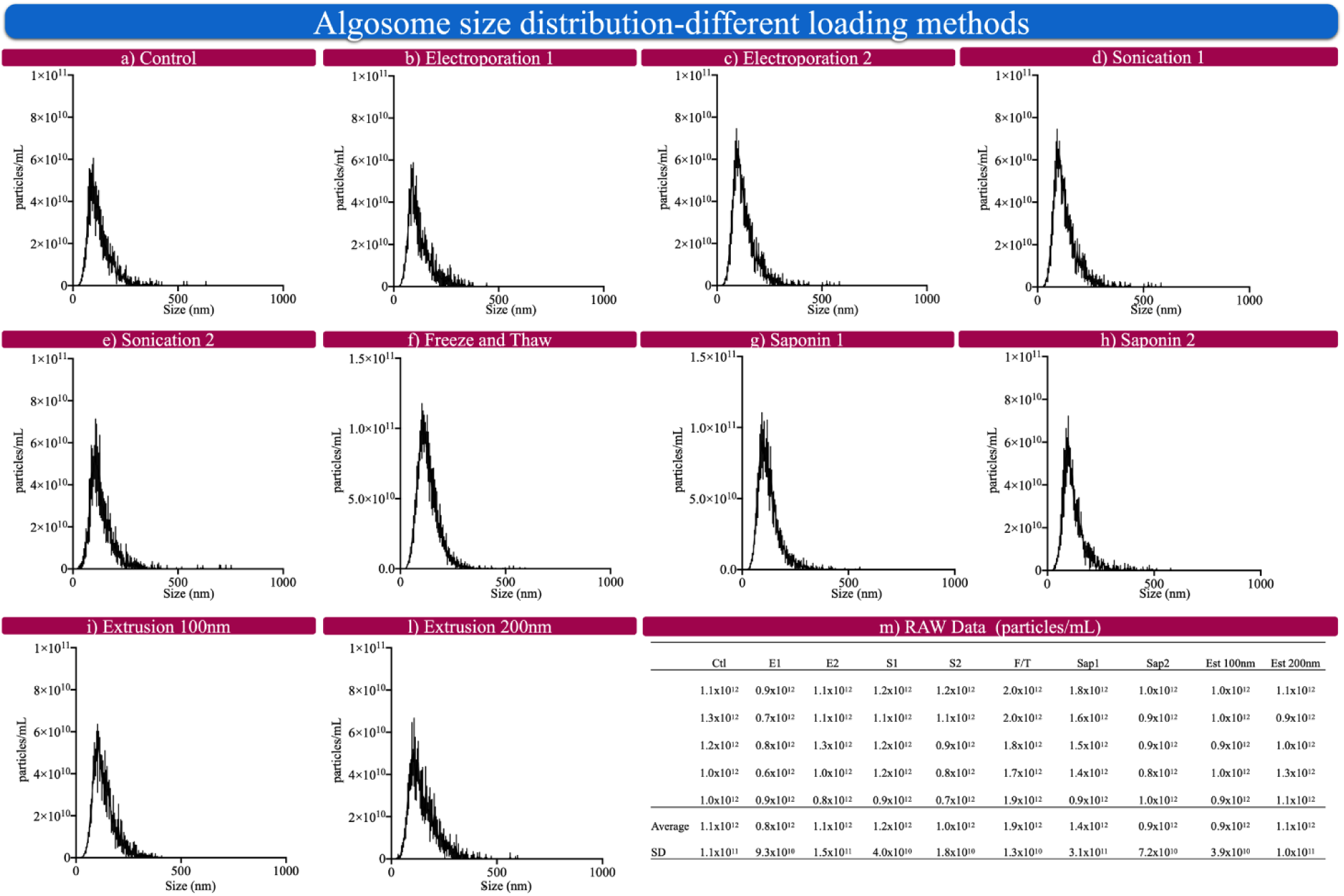
Nanoalgosome size distribution by NTA. a) Untreated nanoalgosomes (*e.g.*, Control) and after different loading methods: b) Electroporation 1; c) Electroporation 2; d) Sonication 1; e) Sonication 2; f) Freeze and Thaw; g) Saponin 1; h) Saponin 2; i) Extrusion 100nm and l) Extrusion 200nm. Each histogram represents the average of five NTA measurements and error bars represent standard deviations. m) Table presents the raw particle count data (particles/mL) for nanoalgosomes stored under different conditions. Each condition was measured in five replicates, and the table provides the particle counts for each replicate. Additionally, the average particle count and standard deviation (SD) for each loading method condition are listed.

## References

1. Adam G, Duncan H: Development of a sensitive and rapid method for the measurement of total microbial activity using fluorescein diacetate (FDA) in a range of soils. Soil Biology and Biochemistry 2001; 33: 943– 951.

2. Adamo G, Fierli D, Romancino DP et al.: Nanoalgosomes: Introducing extracellular vesicles produced by microalgae. Journal of Extracellular Vesicles 2021; 10: e12081.

3. Adamo, G., Santonicola, P., Picciotto, S. et al. Extracellular vesicles from the microalga Tetraselmis chuii are biocompatible and exhibit unique bone tropism along with antioxidant and anti-inflammatory properties. Commun Biol 7, 941 (2024)

4. Bisswanger H: Enzyme assays. Perspectives in Science 2014; 1: 41–55.

5. Bleackley MR, Samuel M, Garcia-Ceron D et al.: Extracellular Vesicles From the Cotton Pathogen Fusarium oxysporum f. sp. vasinfectum Induce a Phytotoxic Response in Plants. Frontiers in Plant Science 2020; 10.

6. Chen C, Sun M, Wang J, Su L, Lin J, Yan X: Active cargo loading into extracellular vesicles: Highlights the heterogeneous encapsulation behaviour. Journal of Extracellular Vesicles 2021; 10: e12163.

7. Chen Y, Zhao Y, Yin Y, Jia X, Mao L: Mechanism of cargo sorting into small extracellular vesicles. Bioengineered 2021; 12: 8186–8201.

8. Choi B, Rempala GA, Kim JK: Beyond the Michaelis-Menten equation: Accurate and efficient estimation of enzyme kinetic parameters. Sci Rep 2017; 7: 17018.

9. Choi D-S, Kim D-K, Choi SJ et al.: Proteomic analysis of outer membrane vesicles derived from Pseudomonas aeruginosa. Proteomics 2011; 11: 3424–3429.

10. Cygler M, Schrag JD, Sussman JL, et al.: Relationship between sequence conservation and three-dimensional structure in a large family of esterases, lipases, and related proteins. Protein Science 1993; 2(3):366–382.

11. de Rond L, van der Pol E, Hau CM et al.: Comparison of Generic Fluorescent Markers for Detection of Extracellular Vesicles by Flow Cytometry. Clinical Chemistry 2018; 64: 680–689.

12. Duggleby RG, Wood C: Analysis of progress curves for enzyme-catalysed reactions. Automatic construction of computer programs for fitting integrated rate equations. Biochem J 1989; 258: 397–402.

13. Duggleby RG: Experimental designs for estimating the kinetic parameters for enzyme-catalysed reactions. J Theor Biol 1979; 81: 671–684.

14. Dzionek A, Dzik J, Wojcieszyńska D, Guzik U: Fluorescein Diacetate Hydrolysis Using the Whole Biofilm as a Sensitive Tool to Evaluate the Physiological State of Immobilized Bacterial Cells. Catalysts 2018; 8: 434.

15. Ender F, Zamzow P, von Bubnoff N, Gieseler F: Detection and Quantification of Extracellular Vesicles via FACS: Membrane Labeling Matters! International Journal of Molecular Sciences 2020; 21: 291.

16. Fontvieille DA, Outaguerouine A, Thevenot DR: Fluorescein diacetate hydrolysis as a measure of microbial activity in aquatic systems: Application to activated sludges. Environmental Technology 1992; 13: 531–540.

17. Fu S, Wang Y, Xia X, Zheng JC: Exosome engineering: Current progress in cargo loading and targeted delivery. NanoImpact 2020; 20: 100261.

18. Gangadaran P, Ahn B-C: Extracellular Vesicle- and Extracellular Vesicle Mimetics-Based Drug Delivery Systems: New Perspectives, Challenges, and Clinical Developments. Pharmaceutics 2020; 12: 442.

19. Garcia-Ceron D, Lowe RGT, McKenna JA et al.: Extracellular Vesicles from Fusarium graminearum Contain Protein Effectors Expressed during Infection of Corn. Journal of Fungi 2021; 7: 977.

20. Gimona M, Brizzi MF, Choo ABH et al.: Critical considerations for the development of potency tests for therapeutic applications of mesenchymal stromal cell-derived small extracellular vesicles. Cytotherapy 2021; 23: 373–380.

21. Gonzales PA, Pisitkun T, Hoffert JD et al.: Large-scale proteomics and phosphoproteomics of urinary exosomes. J Am Soc Nephrol 2009; 20: 363–379.

22. Gonzalez-Begne M, Lu B, Han X et al.: Proteomic analysis of human parotid gland exosomes by multidimensional protein identification technology (MudPIT). J Proteome Res 2009a; 8: 1304–1314.

23. Görgens A, Corso G, Hagey DW et al.: Identification of storage conditions stabilizing extracellular vesicles preparations. Journal of Extracellular Vesicles 2022; 11: e12238.

24. Gray WD, Mitchell AJ, Searles CD: An accurate, precise method for general labeling of extracellular vesicles. MethodsX 2015; 2: 360–367.

25. Han Y, Jones TW, Dutta S et al.: Overview and Update on Methods for Cargo Loading into Extracellular Vesicles. Processes 2021; 9: 356.

26. Herrmann IK, Wood MJA, Fuhrmann G: Extracellular vesicles as a next-generation drug delivery platform. Nat Nanotechnol 2021; 16: 748–759.

27. Hutter JL, Bechhoefer J: Calibration of atomic-force microscope tips. Review of Scientific Instruments 1993; 64: 1868–1873.

28. Jamur MC, Oliver C: Permeabilization of cell membranes. Methods Mol Biol 2010; 588: 63–66.

29. Jeyaram A, Jay SM: Preservation and Storage Stability of Extracellular Vesicles for Therapeutic Applications. AAPS J 2017; 20: 1.

30. Kormelink TG, Arkesteijn GJA, Nauwelaers FA, van den Engh G, Nolte-’t Hoen ENM, Wauben MHM: Prerequisites for the analysis and sorting of extracellular vesicle subpopulations by high-resolution flow cytometry. Cytometry Part A 2016; 89: 135–147.

31. Kusuma GD, Barabadi M, Tan JL, Morton DAV, Frith JE, Lim R: To Protect and to Preserve: Novel Preservation Strategies for Extracellular Vesicles. Frontiers in Pharmacology 2018; 9.

32. LeClaire Michael Gimzewski, James, Sharma Shivani. A review of the biomechanical properties of single extracellular vesicles. Nano Select 2021;2:1–15.

33. Lener T, Gimona M, Aigner L et al.: Applying extracellular vesicles based therapeutics in clinical trials - an ISEV position paper. J Extracell Vesicles 2015; 4: 30087.

34. Liang Y, Lehrich BM, Zheng S, Lu M: Emerging methods in biomarker identification for extracellular vesicle-based liquid biopsy. Journal of Extracellular Vesicles 2021; 10: e12090.

35. Lőrincz ÁM, Timár CI, Marosvári KA et al.: Effect of storage on physical and functional properties of extracellular vesicles derived from neutrophilic granulocytes. J Extracell Vesicles 2014; 3: 10.3402/jev.v3.25465.

36. McMillan HM, Kuehn MJ: Proteomic Profiling Reveals Distinct Bacterial Extracellular Vesicle Subpopulations with Possibly Unique Functionality. Applied and Environmental Microbiology 2022; 89: e01686–22.

37. Michaelis L, Menten ML, Johnson KA, Goody RS: The original Michaelis constant: translation of the 1913 Michaelis-Menten paper. Biochemistry 2011; 50: 8264–8269.

38. Morales-Kastresana A, Telford B, Musich TA et al.: Labeling Extracellular Vesicles for Nanoscale Flow Cytometry. Sci Rep 2017; 7: 1878.

39. Nguyen VVT, Witwer KW, Verhaar MC, Strunk D, van Balkom BWM: Functional assays to assess the therapeutic potential of extracellular vesicles. J Extracell Vesicles 2020; 10: e12033.

40. Nikiforova N, Chumachenko M, Nazarova I et al.: CM-Dil Staining and SEC of Plasma as an Approach to Increase Sensitivity of Extracellular Nanovesicles Quantification by Bead-Assisted Flow Cytometry. Membranes 2021; 11.

41. O’Grady T, Njock M-S, Lion M et al.: Sorting and packaging of RNA into extracellular vesicles shape intracellular transcript levels. BMC Biology 2022; 20: 72.

42. Pachler K, Ketterl N, Desgeorges A et al.: An In Vitro Potency Assay for Monitoring the Immunomodulatory Potential of Stromal Cell-Derived Extracellular Vesicles. Int J Mol Sci 2017; 18: 1413.

43. Paolini L, Monguió-Tortajada M, Costa M et al.: Large-scale production of extracellular vesicles: Report on the “massivEVs” ISEV workshop. Journal of Extracellular Biology 2022; 1: e63.

44. Paterna A, Rao E, Adamo G et al.: Isolation of Extracellular Vesicles From Microalgae: A Renewable and Scalable Bioprocess. Front Bioeng Biotechnol 2022; 10: 836747.

45. Perna RF, Tiosso PC, Sgobi LM et al.: Effects of Triton X-100 and PEG on the Catalytic Properties and Thermal Stability of Lipase from Candida Rugosa Free and Immobilized on Glyoxyl-Agarose. Open Biochem J 2017; 11:66–76.

46. Picciotto S, Barone ME, Fierli D et al.: Isolation of extracellular vesicles from microalgae: towards the production of sustainable and natural nanocarriers of bioactive compounds. Biomater Sci 2021; 9: 2917– 2930.

47. Picciotto S, Santonicola P, Paterna A et al.: Extracellular Vesicles From Microalgae: Uptake Studies in Human Cells and Caenorhabditis elegans. Front Bioeng Biotechnol 2022; 10: 830189.

48. Pocsfalvi G, Turiák L, Ambrosone A et al.: Protein biocargo of citrus fruit-derived vesicles reveals heterogeneous transport and extracellular vesicle populations. J Plant Physiol 2018; 229: 111–121.

49. Pospichalova V, Svoboda J, Dave Z et al.: Simplified protocol for flow cytometry analysis of fluorescently labeled exosomes and microvesicles using dedicated flow cytometer. Journal of Extracellular Vesicles 2015; 4: 25530.

50. Ramirez Marcel I., Amorim Maria G., Gadelha Catarina, Milic Ivana et al.,: Technical challenges of working with extracellular vesicles. Nanoscale 2018; 10(3):881–906.

51. Rankin-Turner S, Vader P, O’Driscoll L, Giebel B, Heaney LM, Davies OG: A call for the standardised reporting of factors affecting the exogenous loading of extracellular vesicles with therapeutic cargos. Advanced Drug Delivery Reviews 2021; 173: 479–491.

52. Raposo G, Stoorvogel W: Extracellular vesicles: exosomes, microvesicles, and friends. Journal of Cell Biology 2013; 200(4):373–83.

53. Ramirez MI, Amorim MG, Gadelha C, et al.: Technical challenges of working with extracellular vesicles. Nanoscale 2018; 10: 881–906.

54. Rizzo J, Chaze T, Miranda K et al.: Characterization of Extracellular Vesicles Produced by Aspergillus fumigatus Protoplasts. mSphere 2020; 5: e00476–20.

55. Robinson PK: Enzymes: principles and biotechnological applications. Essays Biochem 2015; 59: 1–41.

56. Russell AE, Sneider A, Witwer KW, et al. Biological membranes in EV biogenesis, stability, uptake, and cargo transfer: an ISEV position paper arising from the ISEV membranes and EVs workshop. J of Extracellular Vesicle. 2019; 8(1):1684862.

57. Simpson RJ, Kalra H, Mathivanan S: ExoCarta as a resource for exosomal research. Journal of Extracellular Vesicles 2012; 1: 18374.

58. Sivanantham A, Jin Y: Impact of Storage Conditions on EV Integrity/Surface Markers and Cargos. Life 2022; 12: 697.

59. Sırt Çıplak E, Akoğlu KG: Enzymatic Activity as a Measure of Total Microbial Activity on Historical Stone. Heritage 2020; 3: 671–681.

60. Sokolova V, Ludwig A-K, Hornung S et al.: Characterisation of exosomes derived from human cells by nanoparticle tracking analysis and scanning electron microscopy. Colloids Surf B Biointerfaces 2011; 87: 146–150.

61. Sun Y, Liu G, Zhang K, Cao Q, Liu T, Li J: Mesenchymal stem cells-derived exosomes for drug delivery. Stem Cell Res Ther 2021; 12: 561.

62. Sverdlov ED: Amedeo Avogadro’s cry: What is 1 µg of exosomes? BioEssays 2012; 34: 873–875.

63. Tertel T, Schoppet M, Stambouli O, Al-Jipouri A, James PF, Giebel B: Imaging flow cytometry challenges the usefulness of classically used extracellular vesicle labeling dyes and qualifies the novel dye Exoria for the labeling of mesenchymal stromal cell–extracellular vesicle preparations. Cytotherapy 2022; 24: 619–628.

64. USA FDA - Guidance for Industry Potency Tests for Cellular and Gene Therapy Products. Available: https://www.fda.gov/files/vaccines,%20blood%20%26%20biologics/published/Final-Guidance-for-Industry--Potency-Tests-for-Cellular-and-Gene-Therapy-Products.pdf

65. Van Deun J. et al.: EV-TRACK: transparent reporting and centralizing knowledge in extracellular vesicle research. Nat Methods. 2017; 14: 228–232.

66. Van Deun J, Roux Q, Deville S et al.: Feasibility of Mechanical Extrusion to Coat Nanoparticles with Extracellular Vesicle Membranes. Cells 2020; 9: 1797.

67. van de Wakker SI, van Oudheusden J, Mol EA, et al. Influence of short term storage conditions, concentration methods and excipients on extracellular vesicle recovery and function. European Journal of Pharmaceutics and Biopharmaceutics. 2022;170:59–69.

68. Welsh JA, Goberdhan DCI, O’Driscoll L, et al: Minimal information for studies of extracellular vesicles (Misev2023): From basic to advanced approaches. J of Extracellular Vesicle. 2024;13(2):e12404.

69. Yáñez-Mó M, Siljander PR, Andreu Z et al.:Biological properties of extracellular vesicles and their physiological functions. Journal of extracellular vesicles 2015; 4(1):27066.

70. Yekula A, Muralidharan K, Kang KM, Wang L, Balaj L, Carter BS: From laboratory to clinic: Translation of extracellular vesicle based cancer biomarkers. Methods 2020; 177: 58–66.

